# Mimic: A Unified Neural Network Architecture for Modelling Genomic Sequences and Module-level Comparison

**DOI:** 10.64898/2026.10.26.761901

**Authors:** Husam Abdulnabi, J. Timothy Westwood

**Affiliations:** Department of Cell and Systems Biology, University of Toronto Mississauga, Canada

## Abstract

There exist a wide range of Neural Networks (NNs) that have modelled genomic sequences well. The architectures of these NNs consist of three steps: *k*-mer embedding, sequence embedding, and output mapping. The NNs are primarily differentiated from one another by their sequence embedding components. Although different NNs have been compared before, these comparisons are often either limited or involve drastically different NNs. Another issue is that some NNs are configured for a specific sequence length, and different length sequences require manual modification of the architectures. This study examines the architectures of established NNs for genomics sequences. Based on these architectures, a novel NN called Mimic is developed that provides a unified architecture for the principled comparison of NN components.

Mimic was used to model two different types of genomic datasets: protein-DNA interactions and enhancer activity. Along with other architecture components, different sequence embedding strategies were tried such as recurrent layers and convolutional layers. It was found that there exist specific NN components that yield significantly better performing models. Furthermore, Mimic significantly outperformed prevailing standard NNs, Bichrom and DeepSTARR, in modelling protein-DNA interactions and enhancer activity, respectively.

This study provides a framework for more extensive studies toward a general prescription of NN architecture for modelling genomics data.

## 1 Introduction

Neural Networks (NNs) have been widely successful in modelling genomic data. In this study, genomic data refers to nucleotide (nt) sequences and the numeric values (scores) of experimental assays that map to them, referred to as *signals*, which are sequences themselves. Genomic sequences are naturally interpreted as structured relationships among subsequences, or *k*-mers, analogous to the way sentences are composed of words. Accordingly, NNs developed for genomics have often been linked to architectures originally designed for Natural Language Processing.

A broad range of NN architectures has been applied to genomics tasks. Despite their diversity, these models generally share three core components: a *k*-mer embedding step, a sequence embedding step, and an output mapping step. NNs designed for genomics data differ primarily in the components used in the sequence embedding step, reviewed in **Section 2**.

Most genomics tasks involve nt sequences as inputs and signals as outputs, but they often vary substantially in input sequence length. Genomic sequences can range from as few as 100 nts to several hundred thousand nts. Some NNs are typically configured for a fixed sequence length and cannot accommodate variation without manual intervention, which limits their usability and transferability.

In addition to architectural components, each layer contains tunable parameters that strongly influence performance. In practice, only a subset of these hyperparameters is optimized because adjusting certain parameters can produce non-functioning NNs that fail to map inputs to valid outputs. Tuning such parameters often requires manual inspection to ensure proper functioning, which prevents them from being included in automated optimization procedures— including simple methods like random search—thereby limiting overall optimization.

As mentioned, NNs primarily differ in the components used for sequence embedding. While different NNs with varying components have been compared previously, these NNs are typically drastically different and thus conclusions about specific components, especially those concerning sequence embedding, remain weak. To our knowledge, no single unified architecture exists that can execute distinct sequence embedding methods to enable principled comparisons.

Based on prior architectures and guided by implementation and optimization feasibility, **Mimic** was developed: a dynamic and unifying architecture that facilitates both architectural comparison and optimization, detailed in **Section 3**. Mimic automatically adapts to different sequence lengths, supports most sequence embedding methods, and utilizes internal mechanisms for efficient optimization.

Genomics tasks may also involve multiple input sequences. For example, modelling protein–DNA interactions may require both DNA sequences and epigenetic signals such as histone–DNA interactions. Far fewer NN architectures are designed to accommodate multiple input sequences, and doing so introduces additional modelling considerations. We introduce an extension of Mimic designed for multi-sequence inputs called Multi-Mimic in **Section 3.4**.

Mimic is applied to modelling enhancer activity and Multi-Mimic is applied to modelling YY1-genome and CTCF-genome interactions in **Section 4**.

## 2 Review of Genomics Neural Network Architectures

NNs for modelling genomics data have three main steps: *k*-mer embedding, sequencing embedding, and output mapping.

### 2.1 *k*-mer embedding

*k*-mer embedding refers to representing subsequences of length *k* as *v*-dimensional vectors. For a sequence of length *L*, there are *L* − *k* + 1 overlapping *k*-mers, each mapped to a vector, producing an output of shape (*L* − *k* + 1,*v*). One approach derived from Natural Language Processing is to assign each possible *k*-mer a unique learnable vector, a strategy often termed *lookup embedding* (or *token embedding)* (e.g. Ji et al., 2021). However, because the number of possible *k*-mers grows exponentially as 4*^k^*, this method is only feasible for small *k* values. Larger *k* values require tracking and learning embeddings for many *k*-mers, resulting in prohibitively large models and substantial memory requirements.

A more effective strategy is to embed *k*-mers based on their compositional similarity using a convolutional layer, which we refer to here as *compositional embedding*. A convolutional layer is composed of *v* convolutional filters that act analogously to position weight matrices (PWMs). A *k*-mer is scored by each of the convolutional filters, yielding a *v*-dimensional vector. *k*-mers with similar nt composition naturally map to similar vectors. Importantly, convolutional filters can span much larger subsequences (e.g., 20–30 nts or more) while remaining computationally efficient and requiring far fewer parameters than lookup embedding. Consequently, compositional embedding has become the prevailing approach in genomics NNs. The choices of *k* and number of convolutional filters (*v*) are tunable parameters.

Compositional embedding uses a single convolutional layer, thereby embedding *k*-mers by PWM-like filters. PWMs assume positional independence within a *k*-mer, but there is evidence of intra-*k*-mer dependencies that can be higher-order and non-linear (Bulyk et al., 2002). To capture such dependencies while retaining the benefits of compositional embedding, a series of convolutional layers followed by non-linear activation functions can be used.

Two configurations are possible. In the first, the initial convolutional layer uses a filter of length *k*, and subsequent layers use filters of length 1, analogous to dense (linear) layers applied channel-wise. In the second, each convolutional layer uses a base filter length *m* < *k*, with the final layer using a filter of length less than or equal to *m* (**Algorithm 1**). For example, embedding *k*-mers of length 14 with a base filter of 6, convolutional layers with filter lengths of 6, 6, and 4 are used. We refer to this approach as *higher-order compositional embedding*. To our knowledge, no existing NN for genomics employs this strategy for *k*-mer embedding. The base kernel size and the number of convolutional filters to use in each convolutional layer are hyperparameters.

#### Algorithm 1. Filter lengths for higher order compositional embedding

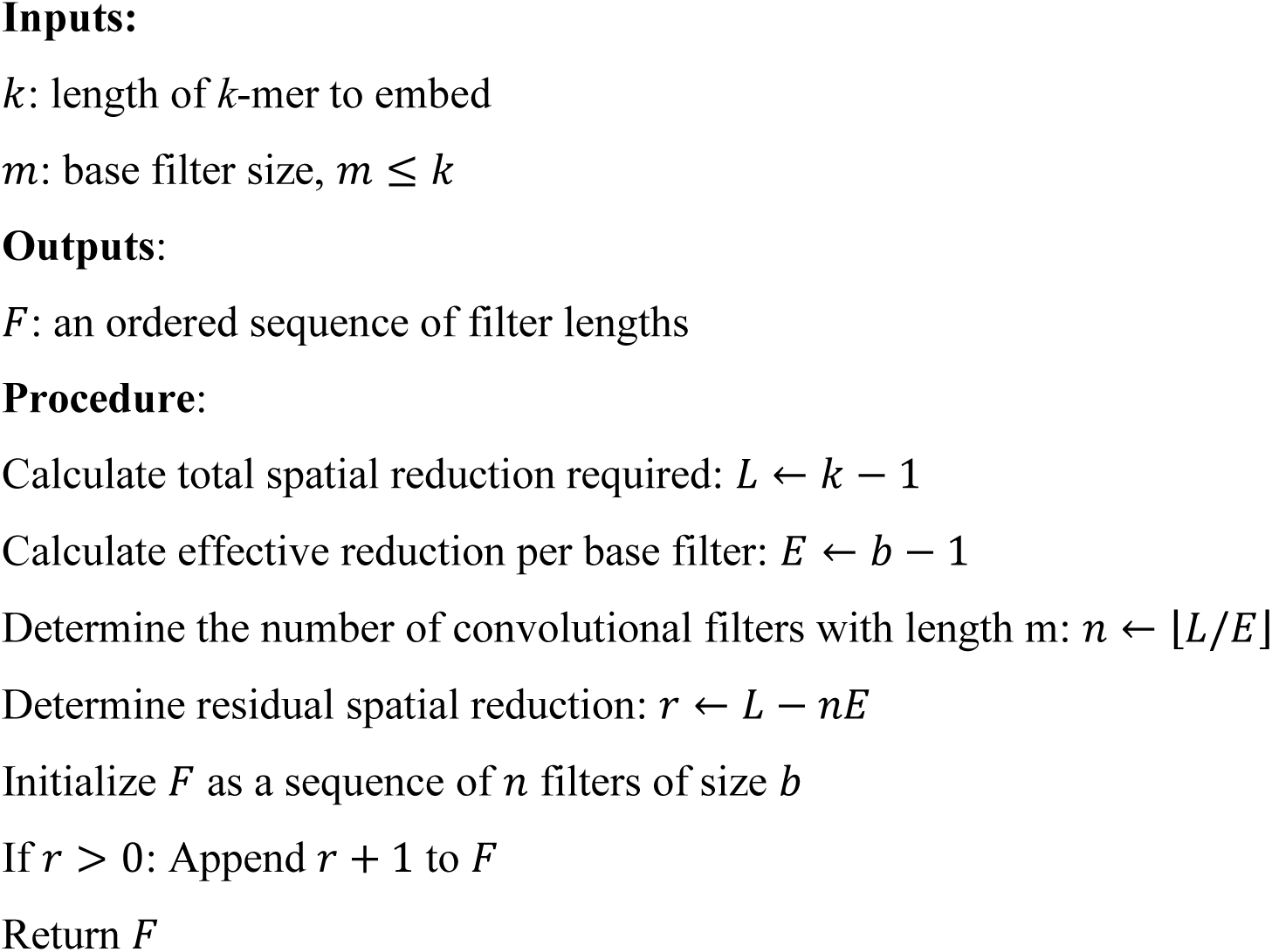

### 2.2 Sequence Embedding

*k*-mer embedding yields an array of (*L* − *k* + 1, *v*). The next step involves summarizing the array of *k*-mer embeddings into a *q*-dimensional vector by considerations of the positions of *k*-mers as well as their relationships with one another, hereafter referred to as *k-mer associations*.

The length of the *k*-mer embeddings can be reduced through a single global pooling layer. Pooling layers such as max and average pooling are devoid of learnable parameters (i.e. non-learnable pooling). Using these pooling layers for global pooling ignores *k*-mer positions and using average pooling further ignores *k*-mer associations. This strategy is employed by DeepBind (Alipanahi et al., 2015) which pools positions by a mixture of average and max pooling. Conversely, global pooling layers can have learnable parameters such as attention-based pooling layers. This approach is used by ti-SFM (Balcı et al., 2023). There are 4 notable attention-based pooling layers: regular, cross-attention (Vaswani et al., 2023), per-dimension (Avsec, Agarwal, et al., 2021), and diagonal (Balcı et al., 2023).

More commonly, local pooling is applied to reduce *k*-mer positions into larger subsequences of length *b* > *k*, referred to as *b*-mers, producing an array of shape (*L* − *b* + 1, *v*). The size of *b* is a hyperparameter.

Attention-based layers such as multi-head attention can be used to modify the *b*-mer embeddings based on the other *b*-mers in the sequence to derive context-specific embeddings, a common strategy in NLP. This approach is employed by AttentiveChrome (Singh et al., 2017), DeepATT (Li et al., 2021), Enformer (Avsec, Agarwal, et al., 2021), ti-SFM (Balcı et al., 2023), DeepCORE (Chandrashekar et al., 2024), and TF-EPI (Liu et al., 2024). The specific parameters of the attention-based layers (e.g. the number of heads for multi-head attention) are additional hyperparameters.

The length of the *b*-mer embeddings can be reduced in several ways. The simplest approach is to use a single dense layer that jointly considers all *b*-mers. The number of nodes of the dense layer is a hyperparameter.

Another approach to reduce *b*-mers is to use recurrent layers which include Long-Short Term Memory (LSTM) and Gated Recurrent Units (GRU) layers. Because recurrent layers are directional, often a second recurrent layer processing in the opposite direction is used in parallel. This approach is used by BiRen (Yang et al., 2017), AttentiveChrome (Singh et al., 2017), DeepATT (Li et al., 2021), BiChrom (Srivastava et al., 2021), and DeepCORE (Chandrashekar et al., 2024). Importantly, recurrent layers process each *b*-mer sequentially and thus can be slow. The number of hidden units within the recurrent layers, the number of layers, and whether the layer is bidirectional are all hyperparameters.

The most widely used approach is to apply a series of modules consisting of a convolutional layer, a non-linear activation function, and a pooling layer. We refer to these modules as Convolutional Pooling Modules (CPMs). Each successive CPM captures longer subsequences and higher-order relationships. This approach effectively captures local and global *k*-mer associations and captures hierarchical relationships. This approach is employed by DeepSEA (Zhou & Troyanskaya, 2015), DanQ (Quang & Xie, 2016), DeepChrome (Singh et al., 2016), Basset (Kelley et al., 2016), AI-TAC (Maslova et al., 2020), DeepSTARR (de Almeida et al., 2022), and TF-EPI (Liu et al., 2024). A similar approach uses modules consisting of a dilated convolutional layer and activation function, employed by Basenji (Kelley et al., 2018), Basenji 2 (Kelley, 2020), and BPNet (Avsec, Weilert, et al., 2021).

CPMs introduce several tunable hyperparameters, including the convolutional filter length, number of filters, pooling kernel length, and pooling stride. These parameters determine the number of modules required, but not all parameter combinations are feasible. For example, a convolutional filter or pooling kernel cannot exceed the length of the incoming sequence. Most NNs hard-code these parameters for a specific sequence length and a specific value of *k*. For different sequence lengths, both the parameters and the number of modules must be manually reconfigured, limiting flexibility and usability.

### 2.3 Output mapping

The output of a NN is of shape (*out_lengt_*_ℎ_, *out_type_*) where *out_type_* refers to different experimental assays. Genomics modelling maps input sequences to other experimental assays which can all vary in length but are all centered on the same nt. While NNs can yield outputs with *out_lengt_*_ℎ_ > 1 (e.g. BPNet) doing so means translational shifts yield overlapping outputs that are potentially inconsistent. This complicates model analysis and final predictions (combining overlapping outputs). This study focuses on *out_lengt_*_ℎ_ of 1.

For *out_lengt_*_ℎ_ of 1, the universal approach to map the sequence embedding to the output involves using 1 or more modules composed of a dense layer, an activation function, and optional dropout layer. The choice of the number of modules, the number of nodes per dense layer, and the amount of dropout are hyperparameters.

## 3 Mimic

While several effective architectural patterns are known, constructing a functioning NN, and subsequently a well-performing one, still requires selecting appropriate hyperparameters. Many combinations of parameters lead to non-functioning NNs, and identifying viable configurations remains a non-trivial component of model design. For NNs that use CPMs that can accept any sequence length and guarantee the appropriate output, we developed automated CPMs. To reduce the number of total parameters to facilitate optimization, we use parameter grouping and number spacing. We combine these elements into a novel NN called Mimic. Furthermore, Mimic can use CPMs for sequence embedding but can also use global pooling, a single dense layer, or recurrent layers, as described in **Section 2**.

We name the model “Mimic” after the Mimic Octopus, which can take on different forms, reflecting the architecture’s ability to adopt multiple sequence embedding mechanisms. Mimic software and a manual with more details are provided in https://github.com/Husam94/poseigen_mimic/.

### 3.1 Automated CPMs

As mentioned, the parameters critical for functioning NNs that use CPMs are the convolutional filter length, pooling kernel length, and pooling stride. Notably, the pooling kernel length must be equal to or greater than the pooling stride. With automated CPMs, the user sets the base convolutional filter length, pooling kernel length, and pooling stride to be used in each CPM and the convolutional filter lengths, pooling kernel lengths, pooling strides, and pooling paddings are automatically determined (**Algorithm 2**). The first CPM does not use a convolutional layer to permit positional reduction of *k*-mer embedding. The components of a CPM are iteratively determined until either convolution or pooling is not possible. This results in the last CPM only consisting of a convolutional layer with filters that span its incoming length.

#### Algorithm 2. Automated convolutional pooling modules

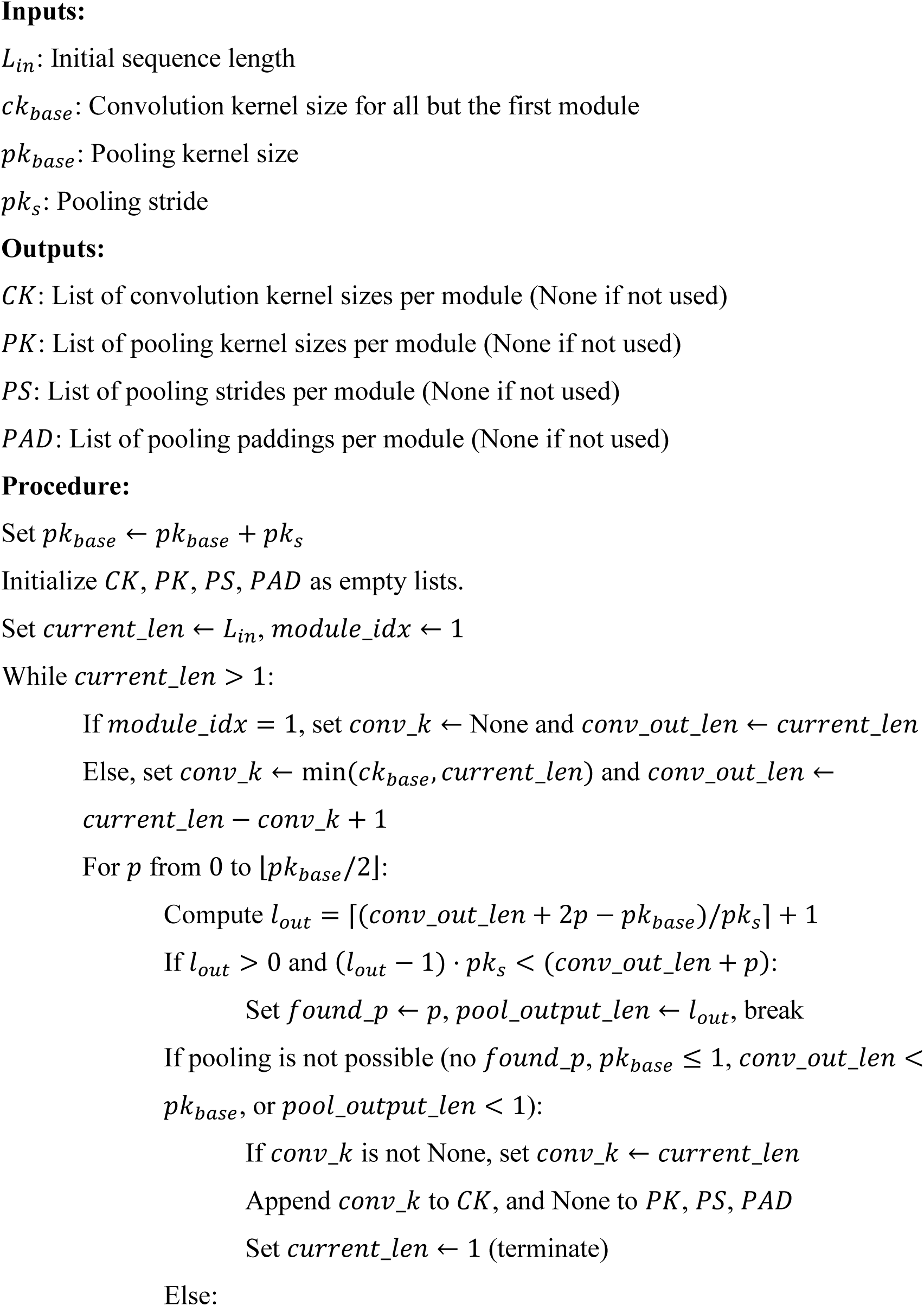
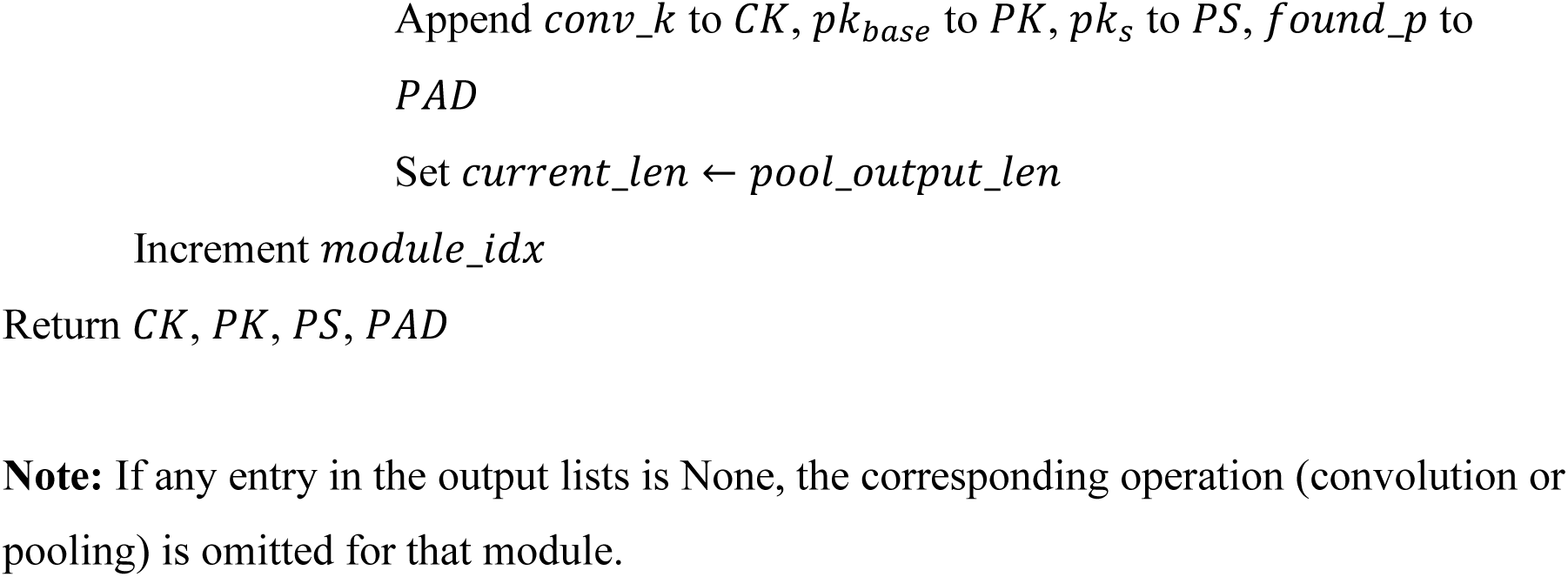

### 3.2 Parameter Grouping and Number Spacing

For higher-order compositional *k*-mer embedding, multiple convolutional layers are used that require specifying the number of filters in each layer. Similarly, for sequence embedding with CPMs, the convolutional layers require specifying the number of filters in each layer. Likewise, for output mapping, multiple dense layers can be used that require specifying the number of nodes in each layer.

Instead of specifying the number of filters of each convolutional layer, we set the number of filters of the last convolutional layer as a multiple of the incoming number of dimensions and apply linear number spacing to impute the number of filters pertaining to the intermediate layers. For example, given an incoming array of 10 dimensions and a multiplier of 4, the final layer will have 40 convolutional filters. Given 3 convolutional layers, the first and second layers will have 20 and 30 convolutional filters, respectively. This method is extended to the number of nodes of the dense layers for output mapping. Since the last dense layer is responsible for the final output, it is automatically set. Instead, the multiplier is for the number of nodes in the penultimate dense layer. This method facilitates the use of different numbers of convolutional filters per convolutional layer and different numbers of nodes per dense layer without setting each one individually.

### 3.3 Mimic

Mimic is a novel NN that can perform higher-order compositional *k*-mer embedding and number spacing for convolutional filters and dense nodes (**Fig. 1**). It can optionally use automated CPMs thereby handling any input sequence length and associated parameters can be thoroughly optimized.

**Fig. 1.**
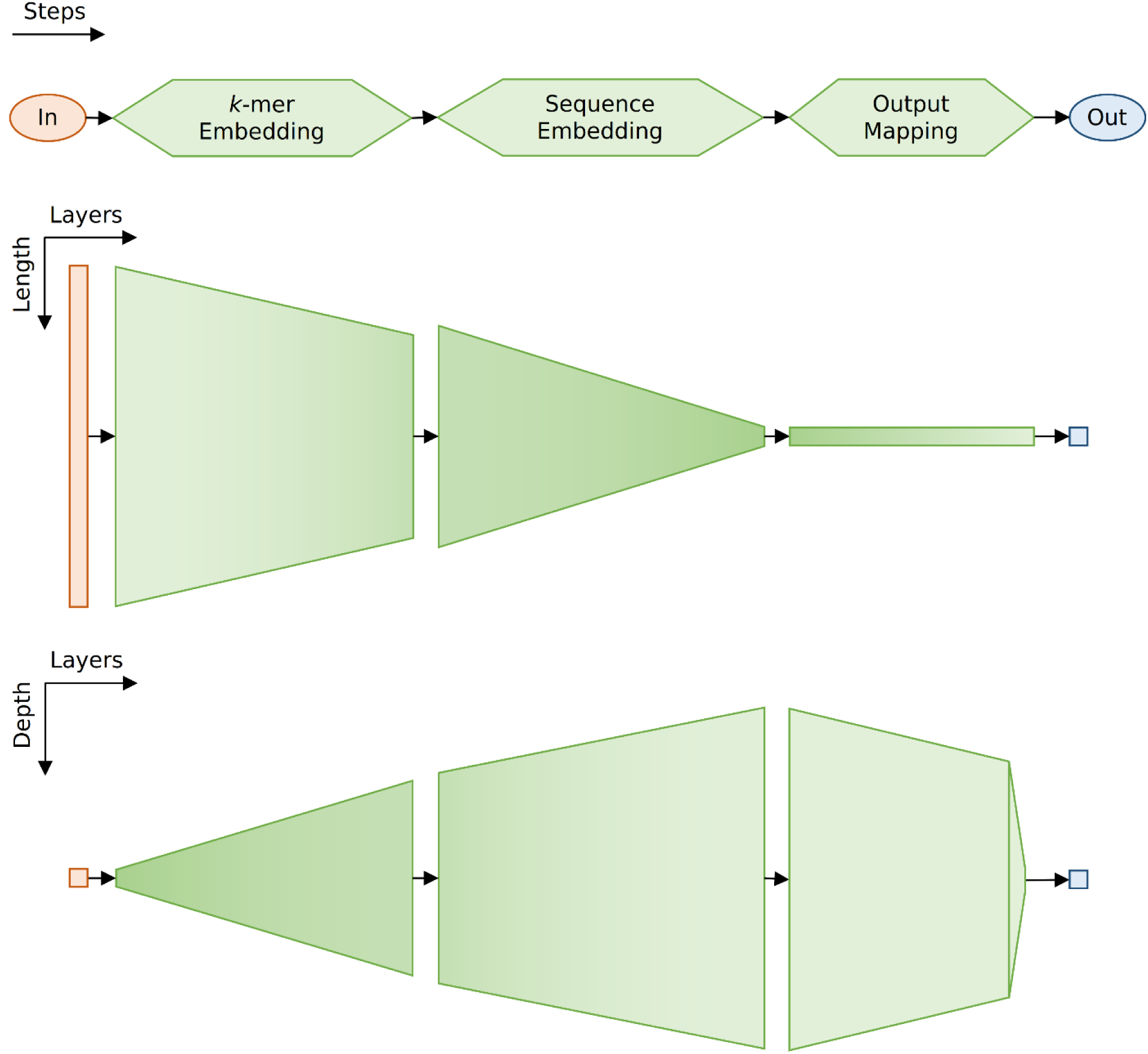
The Mimic model is composed of three steps: *k*-mer embedding, sequence embedding, and output mapping. Diagrams depict different two-dimensional views of an example Mimic model that uses CPMs for sequence embedding with shading representing the missing third dimension. *k*-mer embedding reduces the input sequence length slightly while expanding the embedding dimension (depth). The sequence embedding step reduces the sequence to a single length while further expanding the embedding dimension. The output mapping reduces the embedding dimension slightly to the penultimate layer and then yields the desired output.

Consistent with established NNs for genomics data, Mimic has 3 main steps: *k*-mer embedding, sequence embedding, and output mapping. There are several hyperparameters, all of which can be tuned using automated optimization algorithms (**Text 1**). For *k*-mer embedding, there are 3 hyperparameters: size of *k*, base convolutional filter length, and the multiplier for the number of convolutional filters in the last convolutional layer of the *k*-mer embedding step. For output mapping, there are 2 hyperparameters: the number of dense layers and the multiplier for the number of nodes in the penultimate dense layer of the output mapping step if the number of dense layers > 1. There are also 4 general hyperparameters: the activation function used throughout the model, the final activation function, the amount of dropout on the dense layers of the output mapping step, and whether to add batch normalization after each convolutional and dense layer.

For sequence embedding, Mimic supports several strategies beyond automated CPMs. For sequence embedding with CPMs, there are 4 hyperparameters: the base convolutional filter length, the pooling kernel length, the pooling stride, and the multiplier for the number of convolutional filters in the last convolutional layer of the sequence embedding step. For sequence embedding without CPMs, the size of *b* and the pooling method for reducing *k*-mer embedding are hyperparameters. The size of *b* can be 1, indicating global pooling. If the size of *b* is not 1, a single dense layer can be used. Mimic uses a convolutional layer with filters that span the sequence length instead of a dense layer as they are equivalent but facilitate model design. The number of convolutional filters is a hyperparameter. Alternatively, recurrent layers can be used where the choice of recurrent layer, the number of hidden nodes, the number of layers, and whether to be bidirectional are hyperparameters.

### Text 1: Hyperparameters of the Mimic Model

**Values within square brackets are recommendations for a 500–1500 nt sequence**

#### *k*-mer Embedding

• Size of *k*: [None, 11, 15, 19]

• Base convolutional filter length - can be None, results in a single convolutional layer with filters of size *k*: [None, 5, 7, 9]

• Multiplier for number of convolutional filters in last convolutional layer: [50, 100, 150, 200]

#### Sequence Embedding

- Pooling method: [average, max, *[attention-based methods]]
- Size of *b*: can be None, results in using CPMs: [None, 1, 50, 100]
- If CPM:

`o` Base convolutional filter length: [4, 5, 6]
`o` Pooling stride: [2, 3, 4]
`o` Pooling kernel length: add this to pooling stride to determine length: [0, 1, 2]
- If not CPM:

`o` Layer: [Dense, LSTM, GRU]
`o` If Layer in [LSTM, GRU]:

⯀ Number of layers: [1, 2, 3]
⯀ Bidirectional: [True, False]
- Multiplier for number of convolutional filters in last convolutional layer for CPM, number of convolutional filters in the single dense layer equivalent, or number of hidden units in recurrent layers: [0.5, 1, 1.5, 2]

### Output Mapping

• Number of dense layers: [1, 2, 3, 4]

• Multiplier for number of nodes in penultimate dense layer (if the number of dense layers > 1): [0.5, 1, 1.5, 2]

### General

• Activation Function: [ReLU, Leaky ReLU]

• Final Activation Function: task specific

• Dropout: [0, 0.1, 0.2]

• Batch Normalization: [True, False]

### 3.4 Multi-Mimic

Multi-mimic allows for the integration of multiple different input sequences. This method is useful for some genomics tasks such as modelling protein-DNA interactions that can be dependent on the DNA sequence as well as the local chromatin environment. Input sequences with identical dimensions, such as different histone-DNA sequences of the same length, can be concatenated along the channel dimension and treated as a single sequence.

For each sequence, *k*-mer embedding and sequence embedding are executed by separate modules. There are two options to produce the final output (**Fig. 2**). First, output mapping is executed separately, resulting in multiple Mimic sub-models wherein each sequence is handled separately, referred to as late combination. The outputs of the sub-models are added together, yielding a single output. This strategy is employed by Bichrom (Srivastava et al., 2021). Second, the sequence embeddings are concatenated dimension-wise, yielding a single vector. This single vector is fed to a single joint output mapping step, producing a single output. This method, referred to as early combination, maps cross-interactions of the sequences to the output.

**Fig. 2.**
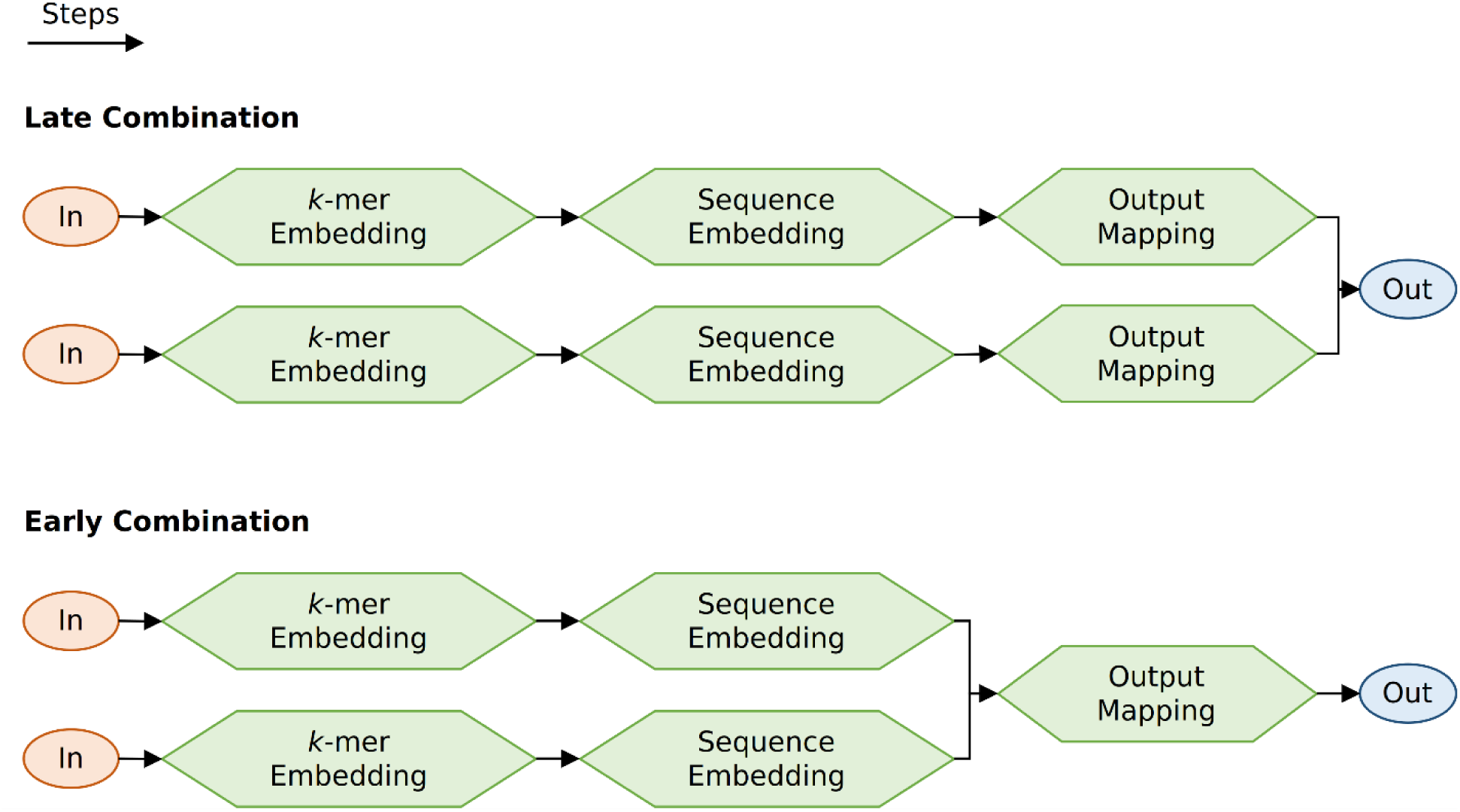
Multi-Mimic processes multiple input sequences. Following Fig. 1, Multi-Mimic can process multiple inputs by either combining the outputs after separate output mapping (late combination) or combining the sequence embeddings for joint output mapping (early combination).

The hyperparameters for each *k*-mer embedding and sequence embedding module are specified separately. Depending on the approach of combining outputs, the hyperparameters for the output mapping module(s) are also specified separately. The approach of combining outputs is an additional hyperparameter. The general hyperparameters (e.g. activation functions) are shared across the whole model, regardless of the combining approach.

## 4 Case Study: Real Biological Datasets

The Mimic model is used on data from two common types of genomics topics: enhancer activity and protein-DNA interactions.

Enhancer activity refers to the varying capabilities of DNA sequences to regulate gene expression. Self-Transcribing Active Regulatory Region (STARR) followed by high throughput sequencing is one of the premier methods to measure enhancer activity (Arnold et al., 2013). With a library of DNA sequences (often fragments of a genome), it involves generating plasmids each containing a DNA sequence downstream of a promoter, transfecting them into cells, and measuring the number of transcribed DNA sequences. A higher transcript count indicates greater positive enhancer activity for that DNA sequence. This technique is especially useful due to its exogenous nature: the enhancer activity is not influenced by chromatin features and is solely based on the DNA sequence and the cellular state. DeepSTARR (de Almeida et al., 2022) is a popular NN used to model enhancer activities, specifically those measured from STARR-seq, and is based on the Basset NN architecture (Kelley et al., 2016). Here, Mimic and DeepSTARR are used to model enhancer activity in a human cancer cell line.

Protein-DNA interactions are principal components of gene regulation and are influenced by the DNA sequence and/or surrounding environment (e.g. histone post-translational modifications, PTMs) (Guertin & Lis, 2010; Slattery et al., 2014; Wang et al., 2012). Yin Yang 1 (YY1) is a DNA sequence-specific transcription factor (a protein that affects gene transcription) implicated in human diseases such as cancer and has a significant role in development, specifically neurogenesis (Verheul et al., 2020). CTCF is also a DNA sequence-specific transcription factor influential in chromatin architecture (Phillips & Corces, 2009). Bichrom is a NN used to model protein-DNA interactions by using DNA sequence and epigenetic signal(s) (Srivastava et al., 2021). Here, the protein-genome interactions of YY1 and of CTCF in a human cancer cell line are modelled using DNA sequence and 11 histone PTMs by Multi-Mimic and Bichrom.

### 4.1 Methods

See https://github.com/Husam94/CS-YY1/, https://github.com/Husam94/CS-CTCF/, https://github.com/Husam94/CS-EnhAct/ for full code on YY1-DNA, CTCF-DNA, and enhancer activity experiments, respectively. Only relevant information is included here.

#### 4.1.1 Data Processing

The datasets for STARR-seq and ChIP-seq in the human cancer cell line K562 were retrieved from the ENCODE database (ENCODE Project Consortium, 2012).

##### 4.1.1.1 Initial

Except for the genome sequence, datasets used in this study are from high-throughput sequencing (HTS) involving a subject and a control. The reads are mapped to the human genome (version hg38), producing read counts per position. The resolution of the read counts was reduced to 50 nt as it did not visually sacrifice specificity. For each subject and control, there are at least 2 biological replicates. HTS reads follow negative binomial distributions. Replicates are scaled by total read counts, producing continuous values that are assumed to follow a gamma distribution. Across replicates, the mean and the sample standard deviation (std) for each value is determined. With so few replicates, the std is shrunk to a smoothed mean-std curve. The mean and shrunken std are used to define the parameters of a gamma distribution.

The control is used to scale the subject values. The ratio of two gamma distributions is a scaled beta prime distribution (SBP, **Eq. 1**) which is equivalent to a generalized beta prime distribution of the second kind with shape parameter *α* = 1 (McDonald & Xu, 1995). For each value, the mode (which is the most probable value from real measurements) is determined from the SBP parameters (**Eq. 2**) and the variance is estimated on a random sample from the SBP. A smoothed mode-variance curve is fit and is used to determine the variance stabilizing transform for the mode (**Eq. 3**, Bartlett, 1947; Love et al., 2014). The variance stabilized modes are the control-scaled values (i.e. intensities) used for modelling.

**Eq. 1. Scaled Beta Prime Distribution**

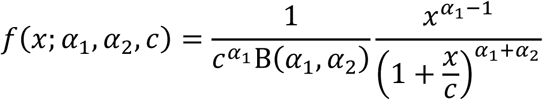

Where:

*x* = control-scaled read count

*α*_1_, *θ*_1_ = shape and scale parameters for the subject gamma distribution

*α*_2_, *θ*_2_ = shape and scale parameters for the control gamma distribution

B(*α*_1_, *α*_2_) = Beta function

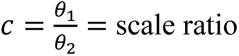

**Eq. 2. Mode of Generalized Beta Prime Distribution**

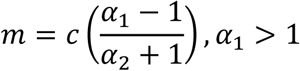

**Eq. 3. Variance Stabilizing Transformation**

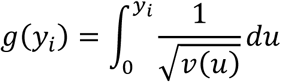

Where:

*y_i_* = is the mode of the SBP at position *i*

*v*(*u*) = variance as a function of the mode, defined by a fitted mode-variance curve

*g*(*y_i_*) = variance-stabilized value

##### 4.1.1.2 Machine Learning Ready Data

The STARR-seq that was performed inserted human genomic DNA sequences of around 800 nts. Accordingly, human genomic DNA sequences of 1000 nts were used to model enhancer activity. Of the possible locations (i.e. sites) in the genome, 30,000 were selected based on specific criteria and sampling (**Sup. Note 10.1.1**). Neighboring windows of ±250 nts and ±500 nts were added, totalling 150,000 observations.

For protein-DNA interactions, human genomic DNA sequences of 500 nts and histone signals of 2000 nts at a 50 nt resolution (signal of length 40) was used. Of the possible locations (i.e. sites) in the genome, 20,000 were selected based on specific criteria and sampling. Neighboring windows of ±150 nts were added, totalling 60,000 observations.

The enhancer activities and protein-DNA interaction intensities are right-skewed (**Fig. 3**). To facilitate modelling, the intensities were centered by the median and scaled by the median-centered absolute max, yielding values that ranged from [-1, 1].

**Fig. 3.**
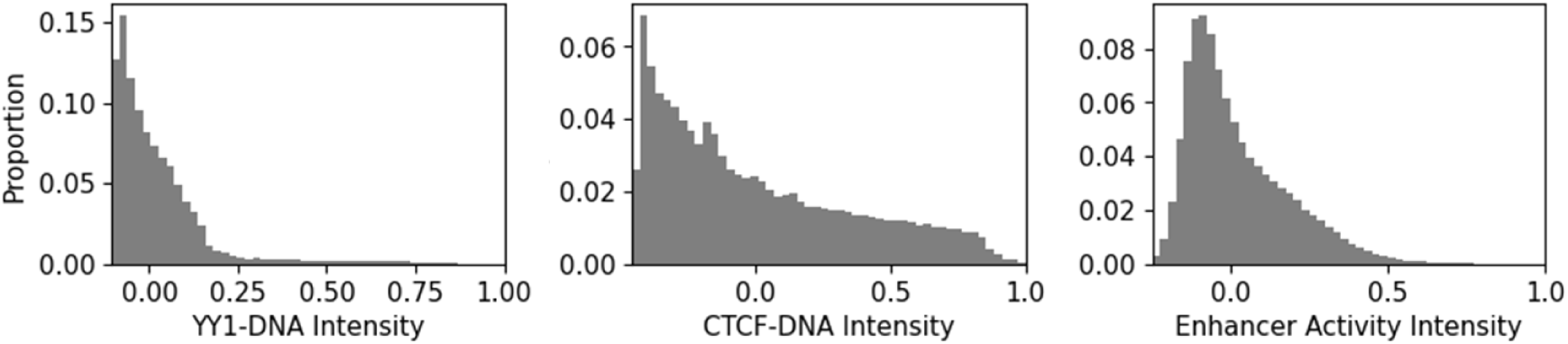
Output values of genomics datasets are right-skewed. Histograms (50 bins) of YY1-DNA intensity (left), CTCF-DNA intensity (right), and enhancer activity intensity (right).

The observations were divided into training, validation, testing, and an additional hold-out set for final evaluation, with proportions of 0.55, 0.15, 0,15, and 0.15, respectively.

#### 4.1.2 Models

All models were fit using the reciprocal-weighted Mean Squared Error as the loss function to account for the right-skewed imbalance (Abdulnabi & Westwood, 2026). Also, all models used in this study used the ADAM optimizer for model fitting (Kingma & Ba, 2014).

##### 4.1.2.1 Mimic Models

Four variants of the Mimic (for enhancer activity) and Multi-Mimic (for protein-DNA interactions) models that differ by sequence embedding method were explored: global attention, single dense layer, recurrent layers, and CPMs. The four variants shared identical search spaces for *k*-mer embedding, output mapping, and global hyperparameters. For CPMs, only average and max pooling were tried as the pooling function as adding learnable pooling methods yields a larger search space and larger models. The architectures of each variant as well as the learning rate and batch size were explored by a random search of 300 candidates.

Optimization of the model architecture in this study involves a two-step procedure. The first step is random search. The second step involves retrieving the top 10 candidates based on the validation set and conducting 10 model repeats for each candidate. The top candidate is then selected based on having the best average score on the validation set.

Model fitting is dependent on stochastic processes, as weight initialization and tuning of weights involve random processes. A single model repeat is an unstable measurement. For robust measurements, the second step of optimization and downstream model fitting involves repeating model fitting. The first step of optimization does not repeat model fitting due to computing budgets and a large search space.

For a robust comparison of model components, the optimization functional of selecting the top 10 candidates was used. The model components were compared by first grouping candidates with a specific component of interest, counting the number of candidates belonging to each group, and determining the expected percentage of top 10 candidates to contain a specific component of interest based on the group sizes. Then, determining the actual percentage of top 10 candidates with the specific component and normalizing it by the expected percentage. This process was repeated on bootstrapped candidates to simulate repeating random search. From the bootstrapped scores, the median, the 5^th^ and 95^th^ percentiles were determined for comparison.

For the top Mimic variant, the top 10 candidates were repeated 10 times. The top candidate was determined as having the best average score on the validation set. The top candidate was repeated an additional 5 times, totaling 15 repeats. For robustness, the repeats were bootstrapped to simulate multiple sets of model repeats. For each set of bootstrapped repeats, the top 3 models (scored on the validation set) were determined and their predictions on the testing set were averaged, akin to an average ensemble. The predictions of the ensemble are scored using reciprocal-density weighted Root Mean Squared Error (recip-RMSE, a lower score is preferred) and reciprocal-density weighted Pearson Correlation Coefficient (recip-PCC, a higher score is preferred). Across bootstrapped average ensembles scores, the median, the 5^th^ and 95^th^ percentiles were determined for comparison.

To visualize final performance, the model indices of the top models across all bootstrapped repeats are used to determine a weighting by scaling the frequency by the total number of model indices. The predictions of the model fitting repeats are combined into a weighted average ensemble.

##### 4.1.2.2 Comparators

TF-Lasso was used as a non-NN control model (de Almeida et al., 2022). For DNA sequences, the human motifs were retrieved from the HOCOMOCO database, specifically from version 11 of the core human motifs (Kulakovskiy et al., 2018) and the TF counts were determined as described in de Almeida et al., 2022.

For histone signals (if applicable), *k*-mers of size 5 (representing 250 nt) across all histone signals were first clustered into 500 clusters by KMeans clustering. The clusters were reduced for redundancy using Agglomerative clustering with a distance cut-off determined by identifying the elbow/knee point by Kneedle algorithm through the *Kneed* python package (Satopaa et al., 2011). Then, histone signals were represented by cluster counts.

The TF counts for DNA sequence and the cluster counts for histone signals (if applicable) were combined into a single vector for each observation. A Lasso model was used and was fit by weighting the samples by the reciprocal-density. The alpha of the Lasso model was optimized through random search of 20 candidates.

For enhancer activity, the DeepSTARR model was used as a NN comparator. The DeepSTARR model originally accepts sequences of 249 nts and performs sequence embedding using 4 CPMs. To handle the roughly four times longer sequence, two optional CPMs were manually added based on their last CPM. The number of CPMs (4, 5, or 6) was an additional hyperparameter, along with their other hyperparameters. The DeepSTARR model was optimized through the same two-step procedure as the Mimic model.

For protein-DNA interactions, the Bichrom model was used as a NN comparator. The Bichrom model originally accepts nucleotide sequences and histone signals matching that used in this study so no additional modifications were done. The Bichrom model was optimized through the same two-step procedure as the Mimic model.

### 4.2 Results and Discussion

For simplicity, Mimic and Multi-Mimic are referred to as simply Mimic in this section.

#### 4.2.1 Mimic Components

For Mimics that use global attention-based pooling, the 4 attention-based pooling methods mentioned earlier were tried (**Table 1**). For YY1-DNA interactions, models with per-dimension attention pooling layer perform the best albeit statistically similar to models with the other attention-based pooling layers. For CTCF-DNA and enhancer activity, however, models with regular and cross-attention pooling layers perform the best albeit statistically similar to models with other attention-based pooling layers.

**Table 1.** Mimics that use global attention-based pooling with cross-attention layers perform as well if not better than those with per-dimension and diagonal attention layers. The median (“Med”), 5^th^ percentile (“p5”), and 95^th^ percentile (“p95”) of bootstrap percent top 10 scores across datasets. A greater score is preferred. “PerDim” refers to per-dimension attention layer.

|  | YY1-DNA |  |  | CTCF-DNA |  |  | Enhancer Activity |  |  |
| --- | --- | --- | --- | --- | --- | --- | --- | --- | --- |
|  | Med | p5 | p95 | Med | p5 | p95 | Med | p5 | p95 |
| PerDim | 21.9 | -38.8 | 92.3 | -100.0 | -100.0 | -8.0 | -29.0 | -78.9 | 36.4 |
| Diagonal | 9.1 | -59.5 | 81.9 | -49.0 | -100.0 | 5.6 | -47.1 | -83.9 | 7.2 |
| Regular | -27.0 | -77.7 | 29.6 | 40.0 | -24.1 | 109.4 | 38.2 | -25.9 | 111.0 |
| Cross | -2.6 | -63.0 | 71.4 | 66.7 | -4.3 | 145.9 | 40.6 | -26.3 | 117.8 |

For Mimics that use recurrent layers, 2 recurrent layers (GRU and LSTM) were tried with different combinations of number of layers and bidirectionality (**Table 2**). Models that used GRU layers performed as well as, if not statistically better than those that used LSTM layers. Models using 2 recurrent layers performed best for modelling YY1-DNA and CTCF-DNA interactions but models using 3 recurrent layers performed best for modelling enhancer activity. Moreover, models that used unidirectional recurrent layers performed better (albeit statistically similar) to models that used bidirectional recurrent layers across all datasets.

**Table 2.** Mimics that use recurrent layers with GRU layers outperform those with LSTM layers. “Uni” and “Bi” refer to unidirectional and bidirectional recurrent layers, respectively. See Table 1 for additional descriptions.

|  | YY1-DNA |  |  | CTCF-DNA |  |  | Enhancer Activity |  |  |
| --- | --- | --- | --- | --- | --- | --- | --- | --- | --- |
| Component | Med | p5 | p95 | Med | p5 | p95 | Med | p5 | p95 |
| GRU | 29.6 | -4.5 | 64.5 | 9.8 | -30.5 | 41.9 | 73.2 | 35.4 | 106.1 |
| LSTM | -34.3 | -68.8 | 5.0 | -10.1 | -43.4 | 30.5 | -69.1 | -91.1 | -33.5 |
| 1 Layer | -0.9 | -45.0 | 45.2 | -21.4 | -70.6 | 28.6 | -47.8 | -85.7 | -0.9 |
| 2 Layers | 33.3 | -16.7 | 89.2 | 77.4 | 19.3 | 139.4 | -17.0 | -58.7 | 32.8 |
| 3 Layers | -40.0 | -83.9 | 9.4 | -55.0 | -85.7 | -7.2 | 70.1 | 13.2 | 125.8 |
| Uni | 7.1 | -29.1 | 44.8 | 27.5 | -17.3 | 66.7 | 10.4 | -29.1 | 41.5 |
| Bi | -7.5 | -44.1 | 29.5 | -23.9 | -60.8 | 15.4 | -11.2 | -43.3 | 28.6 |

Across all Mimic variants, Mimics that conduct higher-order compositional *k*-mer embedding for DNA sequences performed statistically as well if not better than those that conduct compositional *k*-mer embedding (**Table 3**). For modelling YY1-DNA interactions, models that conduct compositional *k*-mer embedding of histone signals performed the best albeit statistically similar to the other models. In contrast, for modelling CTCF-DNA interactions, the opposite is true: models that conduct higher order compositional *k*-mer embedding of histone signals performed the best albeit statistically similar to other models. For modelling enhancer activity, models that conduct higher-order compositional *k*-mer embedding with base filter sizes of 7 performed significantly better than models that conduct compositional *k*-mer embedding. Furthermore, for modelling protein-DNA interactions, models with early combination performed statistically as well if not better than models with late combination (**Table 4**).

**Table 3.** Mimics that conduct higher-order compositional *k*-mer embedding for DNA sequences outperform those that conduct compositional *k*-mer embedding. “Comp. Embed” refers to compositional *k*-mer embedding. “Higher Embed (*x*)” refers to higher-order compositional *k*-mer embedding with the base convolutional filter size *x*. “His.” refers to histone. “NA” refers to not applicable. See Table 1 for additional descriptions.

| Component | YY1-DNA |  |  | CTCF-DNA |  |  | Enhancer Activity |  |  |
| --- | --- | --- | --- | --- | --- | --- | --- | --- | --- |
|  | Med | p5 | p95 | Med | p5 | p95 | Med | p5 | p95 |
| DNA Comp. Embed | -70.1 | -100.0 | -35.9 | 5.5 | -44.1 | 58.7 | -71.2 | -100.0 | -35.8 |
| DNA Higher Embed. (7) | 28.0 | -27.3 | 88.6 | -15.3 | -58.1 | 34.8 | 97.3 | 39.1 | 147.9 |
| DNA Higher Embed. (9) | 40.0 | -13.9 | 96.5 | 7.7 | -43.8 | 63.5 | -20.5 | -59.2 | 27.1 |
| His. Comp. Embed | 38.6 | -34.6 | 118.0 | -41.0 | -100.0 | 29.7 | NA | NA | NA |
| His. Higher Embed. (5) | -42.1 | -81.9 | 20.8 | 17.3 | -42.7 | 92.9 | NA | NA | NA |
| His. Higher Embed. (7) | 4.3 | -60.4 | 81.2 | 3.3 | -59.6 | 83.7 | NA | NA | NA |
| His. Higher Embed. (9) | -4.9 | -62.4 | 66.7 | 13.5 | -54.5 | 77.6 | NA | NA | NA |

**Table 4.** Multi-Mimics that conduct early combination perform as well if not better than models that conduct late combination. “Comb.” refers to combination See Table 1 for additional descriptions.

|  | YY1-DNA |  |  | CTCF-DNA |  |  |
| --- | --- | --- | --- | --- | --- | --- |
| Component | Med | p5 | p95 | Med | p5 | p95 |
| Early Comb. | 11.5 | -27.2 | 47.1 | 30.7 | -12.1 | 65.9 |
| Late Comb. | -11.3 | -48.7 | 25.2 | -30.3 | -69.2 | 12.6 |

Lastly, but arguably most importantly, across all Mimic variants and all datasets, Mimics that use CPMs significantly outperform all other variants (**Table 5**). Notably, Mimics that use CPMs were limited to non-learnable pooling layers as trying learnable pooling further expands the large search space and model sizes. It may be that using learnable pooling within CPMs leads to further performance benefits and it is recommended to be investigated in future studies.

**Table 5.** CPM Mimic models outperform other Mimic variants. “Global Att.” refers to Global Attention. See Table 1 for additional descriptions.

|  | YY1-DNA |  |  | CTCF-DNA |  |  | Enhancer Activity |  |  |
| --- | --- | --- | --- | --- | --- | --- | --- | --- | --- |
| Component | Med | p5 | p95 | Med | p5 | p95 | Med | p5 | p95 |
| Global Att. | -100.0 | -100.0 | -100.0 | -100.0 | -100.0 | -100.0 | -100.0 | -100.0 | -100.0 |
| Single Dense | -100.0 | -100.0 | -57.7 | -100.0 | -100.0 | -100.0 | -38.5 | -81.0 | 34.3 |
| Recurrent | -100.0 | -100.0 | -100.0 | -100.0 | -100.0 | -100.0 | -100.0 | -100.0 | -80.0 |
| CPM | 287.1 | 240.9 | 325.6 | 300.0 | 270.3 | 333.2 | 230.1 | 160.0 | 290.4 |

In general, these findings suggest that higher-order compositional *k*-mer embedding is most effective for *k*-mer embedding and that CPMs are the most effective for sequence embedding. Still, these findings are limited for several reasons. First, although the datasets span two important genomics topics, they do not span all genomics topics and this study only used three datasets. Second, the search space was curated based on computational budgets and previous results (not included here). It may be that expanding the search space (e.g. using recurrent layers with 4 layers) yields different results. Third, although 300 candidates were tried for each variant of Mimic (limited by computational budgets), this may not be sufficiently exhaustive of the search space: more candidates may yield different results.

Nonetheless, this study exemplifies the utility of the Mimic model to investigate the efficacy of different model components. It is recommended that similar trials are conducted for different datasets.

#### 4.2.2 Final Performance

Mimic models significantly outperformed TF-Lasso and other NN models across datasets measured by recip-RMSE (**Table 6**) and recip-PCC (**Table 7**). Measured by recip-RMSE, on YY1-DNA and CTCF-DNA interactions, Mimic outperformed Bichrom by 12.3% and 13.4%, respectively. On enhancer activity, Mimic outperformed DeepSTARR by 11.4%.

**Table 6.**
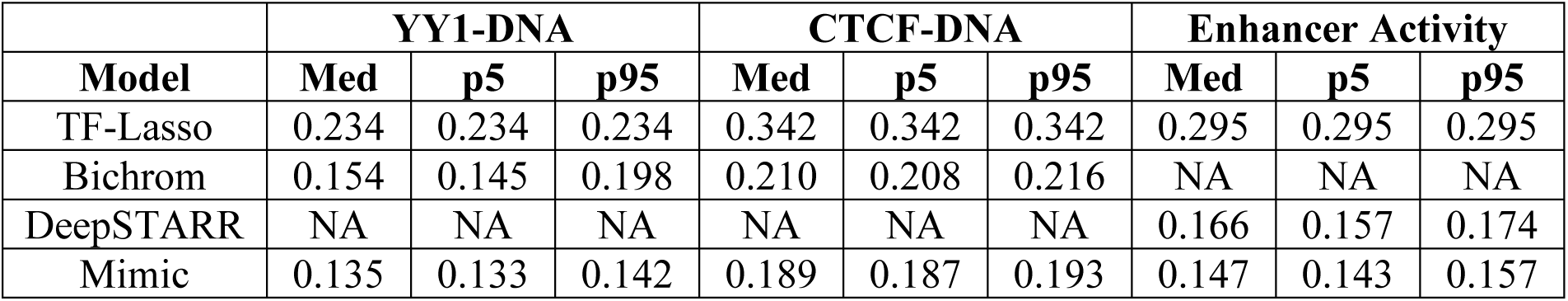
Mimic model outperforms other models across datasets measured by recip-RMSE. The median (“Med”), 5^th^ percentile (“p5”), and 95^th^ percentile (“p95”) of recip-RMSE scores of bootstrapped average ensemble predictions on the testing set. A lower score is preferred. “NA” refers to not applicable.

|  | YY1-DNA |  |  | CTCF-DNA |  |  | Enhancer Activity |  |  |
| --- | --- | --- | --- | --- | --- | --- | --- | --- | --- |
| Model | Med | p5 | p95 | Med | p5 | p95 | Med | p5 | p95 |
| TF-Lasso | 0.234 | 0.234 | 0.234 | 0.342 | 0.342 | 0.342 | 0.295 | 0.295 | 0.295 |
| Bichrom | 0.154 | 0.145 | 0.198 | 0.210 | 0.208 | 0.216 | NA | NA | NA |
| DeepSTARR | NA | NA | NA | NA | NA | NA | 0.166 | 0.157 | 0.174 |
| Mimic | 0.135 | 0.133 | 0.142 | 0.189 | 0.187 | 0.193 | 0.147 | 0.143 | 0.157 |

**Table 7.**
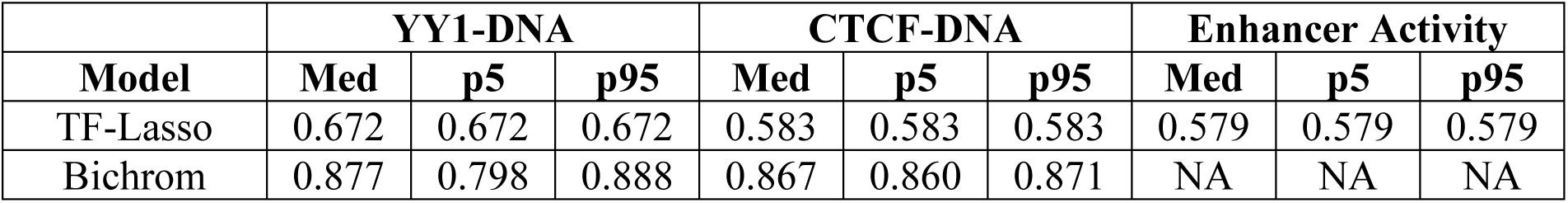
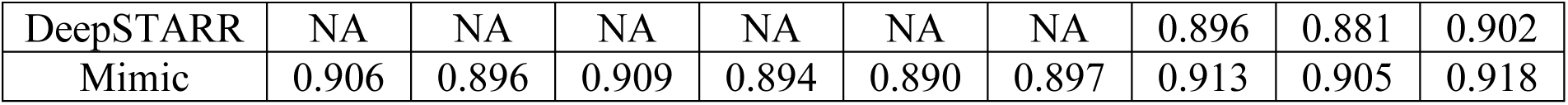
Mimic model outperforms other models across datasets measured by recip-PCC. The median, 5^th^ percentile, and 95^th^ percentile of recip-PCC scores of bootstrapped average ensemble predictions on the testing set. A higher score is preferred. “NA” refers to not applicable.

|  | YY1-DNA |  |  | CTCF-DNA |  |  | Enhancer Activity |  |  |
| --- | --- | --- | --- | --- | --- | --- | --- | --- | --- |
| Model | Med | p5 | p95 | Med | p5 | p95 | Med | p5 | p95 |
| TF-Lasso | 0.672 | 0.672 | 0.672 | 0.583 | 0.583 | 0.583 | 0.579 | 0.579 | 0.579 |
| Bichrom | 0.877 | 0.798 | 0.888 | 0.867 | 0.860 | 0.871 | NA | NA | NA |
| DeepSTARR | NA | NA | NA | NA | NA | NA | 0.896 | 0.881 | 0.902 |
| Mimic | 0.906 | 0.896 | 0.909 | 0.894 | 0.890 | 0.897 | 0.913 | 0.905 | 0.918 |

Measured by recip-PCC, on YY1-DNA and CTCF-DNA interactions, Mimic outperformed Bichrom by 3.2% and 3.0%, respectively. On enhancer activity, Mimic outperformed DeepSTARR by 1.9%. The predictions of weighted ensembles of NN models are visually similar (**Fig. 4**).

**Fig. 4.**
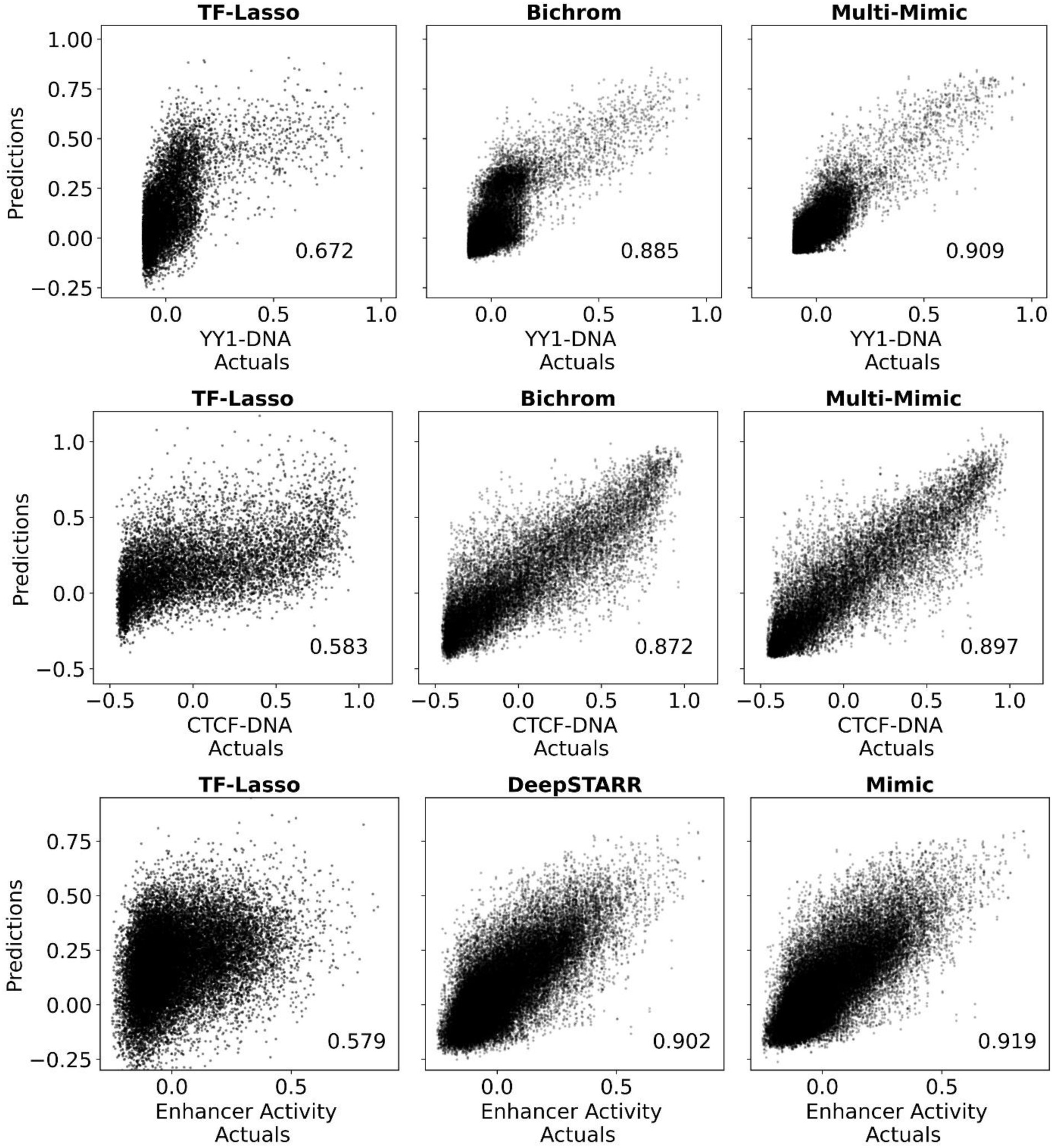
Scatterplots of weighted ensemble predictions of different models on different datasets. The predictions and actuals pertain to the testing set of respective datasets. The recip-PCC is included in each plot at the bottom right.

## 5 Future Directions

The Mimic framework establishes a foundation for systematic architectural comparison, but several avenues remain for further investigation to improve the optimization and representational capacity of genomic NNs. There are four main ones described here.

First, the current implementation uses linear number spacing, but geometric number spacing provides a promising alternative. The current Mimic implementation allows for geometric number spacing. Preliminary trials outside this study showed that geometric number spacing does not improve performance further. Also, geometric number spacing introduces another variable for optimization (the geometric ratio). Still, more trials are warranted.

Second, the use of self-attention for context aware *k*- or *b*-mer embedding may improve performance. Although it was acknowledged as a strategy used in other NNs, it was not benchmarked in this study due to computing resource constraints. Investigating the efficacy of self-attention layers (not to be confused with attention-based pooling) introduces another variable to an already large space and increases model size thereby requiring greater computing resources per model.

Third, dilated convolutional layers may serve as an effective alternative to CPMs as dilated filters have larger receptive fields. However, it is expected that CPMs are more effective as dilated filters are susceptible to the gridding effect where the sparse sampling of sequence can result in the loss of critical local information. CPMs, in contrast, ensure that every nucleotide is accounted for.

Fourth, sequence embedding methods were described and implemented in isolation, but a combination of sequence embedding methods may provide performance benefits. For example, sequence embedding with CPMs followed by recurrent layers, as employed by Min et al., 2017.

## 6 Conclusion

Mimic is a novel NN for modelling genomic sequences with a flexible architecture and internal mechanisms that guarantee functioning NNs. The architecture design distills established NNs for modelling genomics sequences. It enables principled comparisons of NN components using a single unified architecture and facilitates optimization.

Mimic was applied on modelling protein-DNA interactions and enhancer activity in a human cancer cell line. It was found that Mimic models that consist of higher-order compositional *k*-mer embedding for DNA sequences and those that use CPMs for sequence embedding perform as well if not better than any other Mimic variants. These findings suggest that higher-order compositional *k*-mer embedding for DNA sequences and CPMs for sequence embedding be used for modelling genomics sequences. These findings, however, are limited and it is suggested that similar trials are conducted for other datasets. Furthermore, Mimic significantly outperformed popular NNs of Bichrom on modelling protein-DNA interactions and DeepSTARR on modelling enhancer activity.

## 7 Acknowledgements

None.

## 8 Author Contributions

**Conceptualization:** Husam Abdulnabi

**Data curation:** Husam Abdulnabi

**Formal analysis:** Husam Abdulnabi

**Investigation:** Husam Abdulnabi

**Methodology:** Husam Abdulnabi

**Project administration:** Husam Abdulnabi

**Software:** Husam Abdulnabi

**Supervision:** J. Timothy Westwood

**Writing – original draft:** Husam Abdulnabi

**Writing – review & editing:** J. Timothy Westwood

## 10 Supplementary

### 10.1 Notes

#### 10.1.1 Site Selection

Genomics experiments often form peaks which signify enrichment. Peak calling with MACS was used to determine peak sizes of targets (the outputs for modelling). Based on the peak size, the greatest local values (flags), were determined such that no two flags are within peak size of one another. Along with the flags, the neighboring values are used as translational augments.

The flags were filtered for two criteria. First, each flag and its neighboring values must not be within Blacklist regions. Second, each flag and its neighboring values must have equal or greater than the 5^th^ percentile of mean control read values that are greater than 0 to ensure adequate number of control reads.

Of the remaining flags, a desired number of flags were selected so that the distribution of flag values was as balanced as possible by a method similar to weighted random sampling with reciprocal-density weights and without replacement.

### 10.2 Texts

**Sup. Text. 1.**
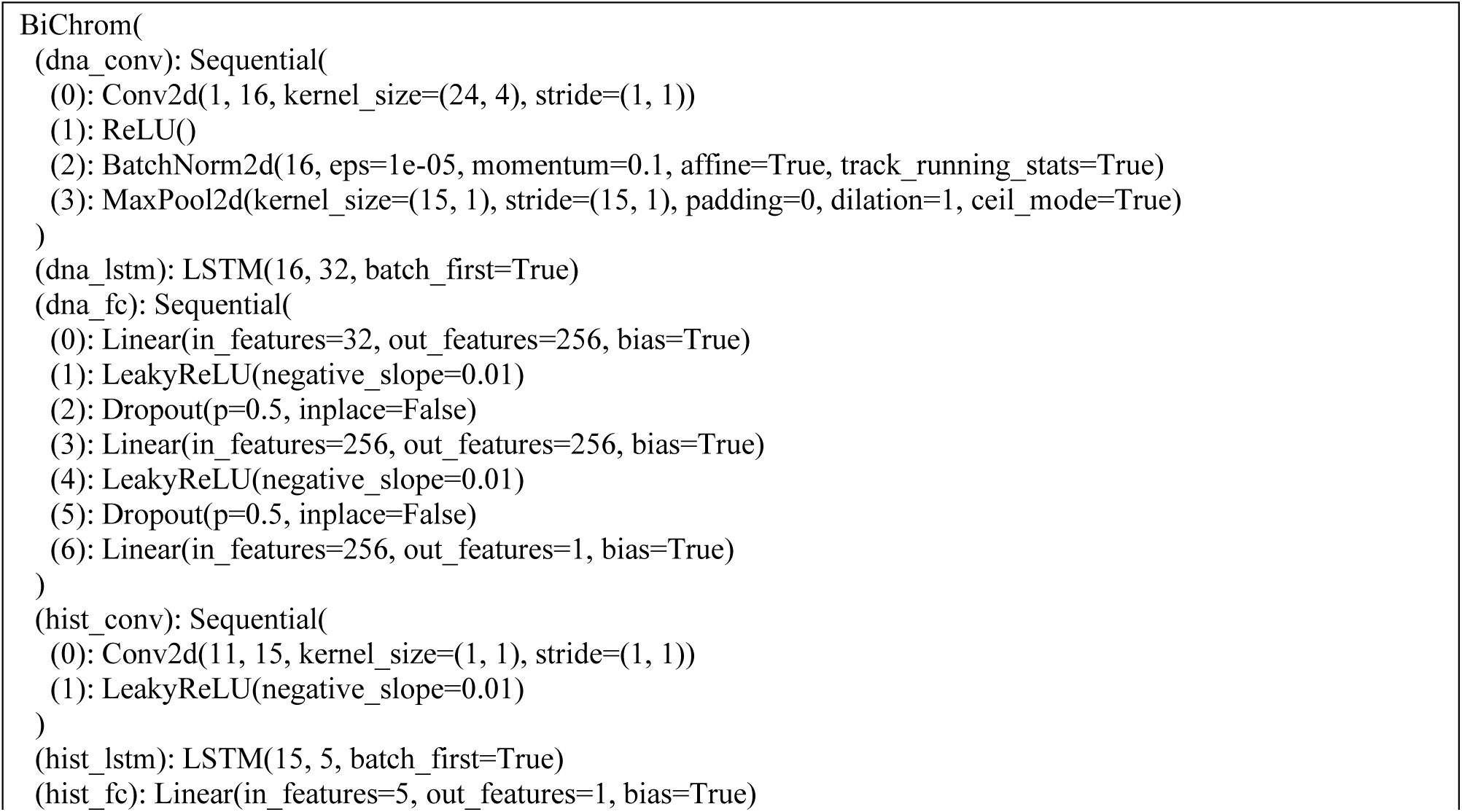
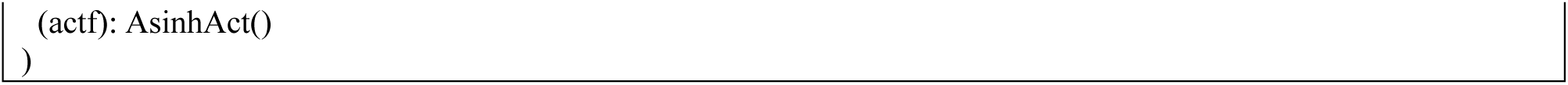
Optimized Bichrom architecture for modelling YY1-DNA interactions.

**Sup. Text. 2.**
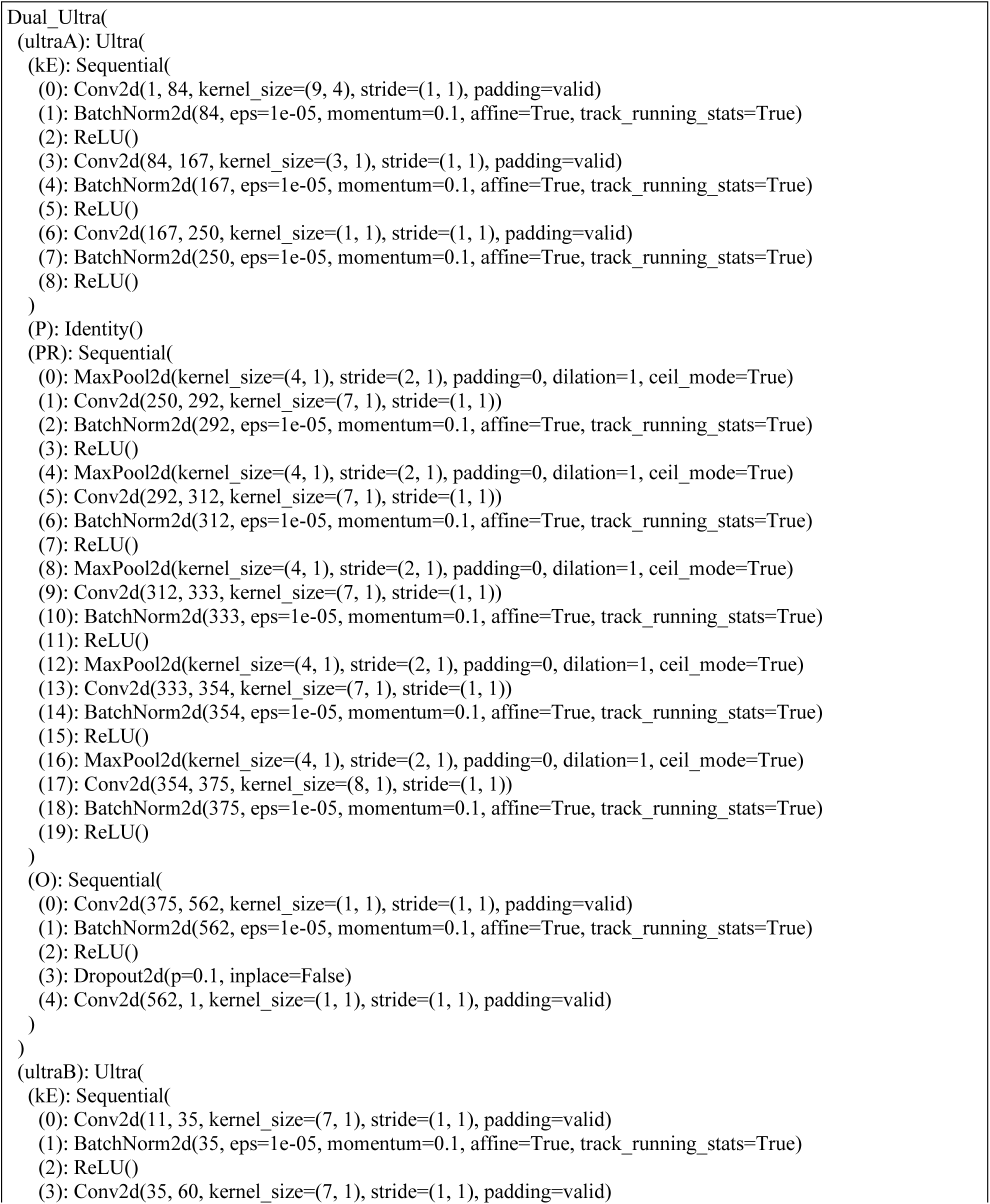
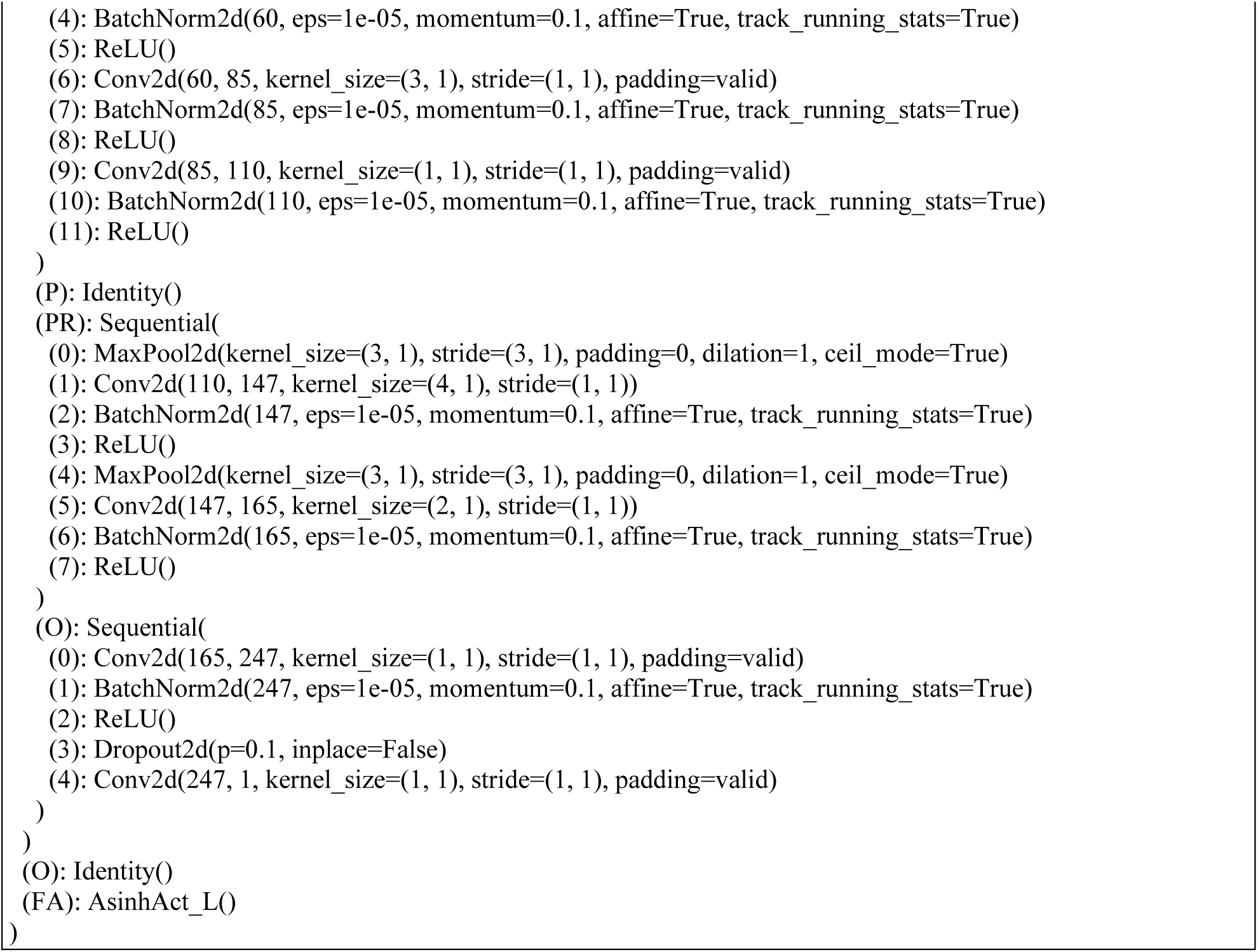
Optimized Mimic architecture for modelling YY1-DNA interactions.

**Sup. Text. 3.**
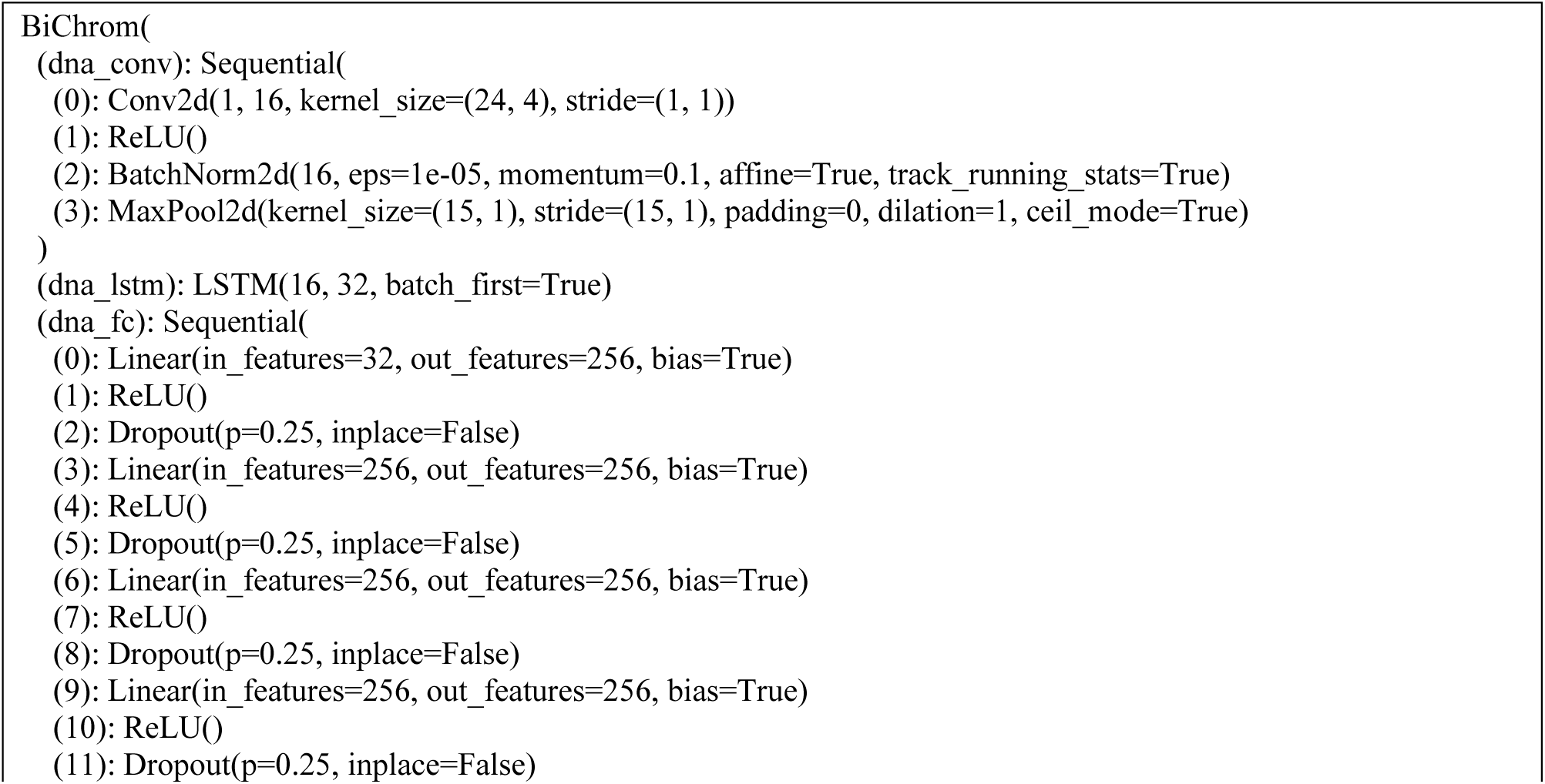
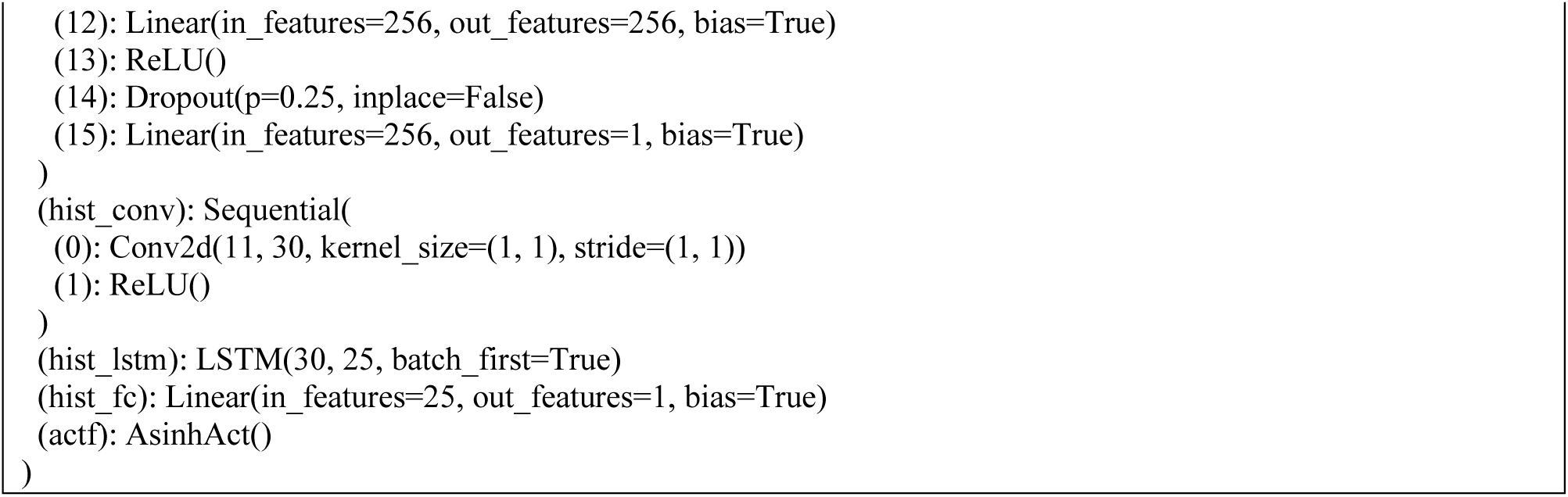
Optimized BiChrom architecture for modelling CTCF-DNA interactions.

**Sup. Text. 4.**
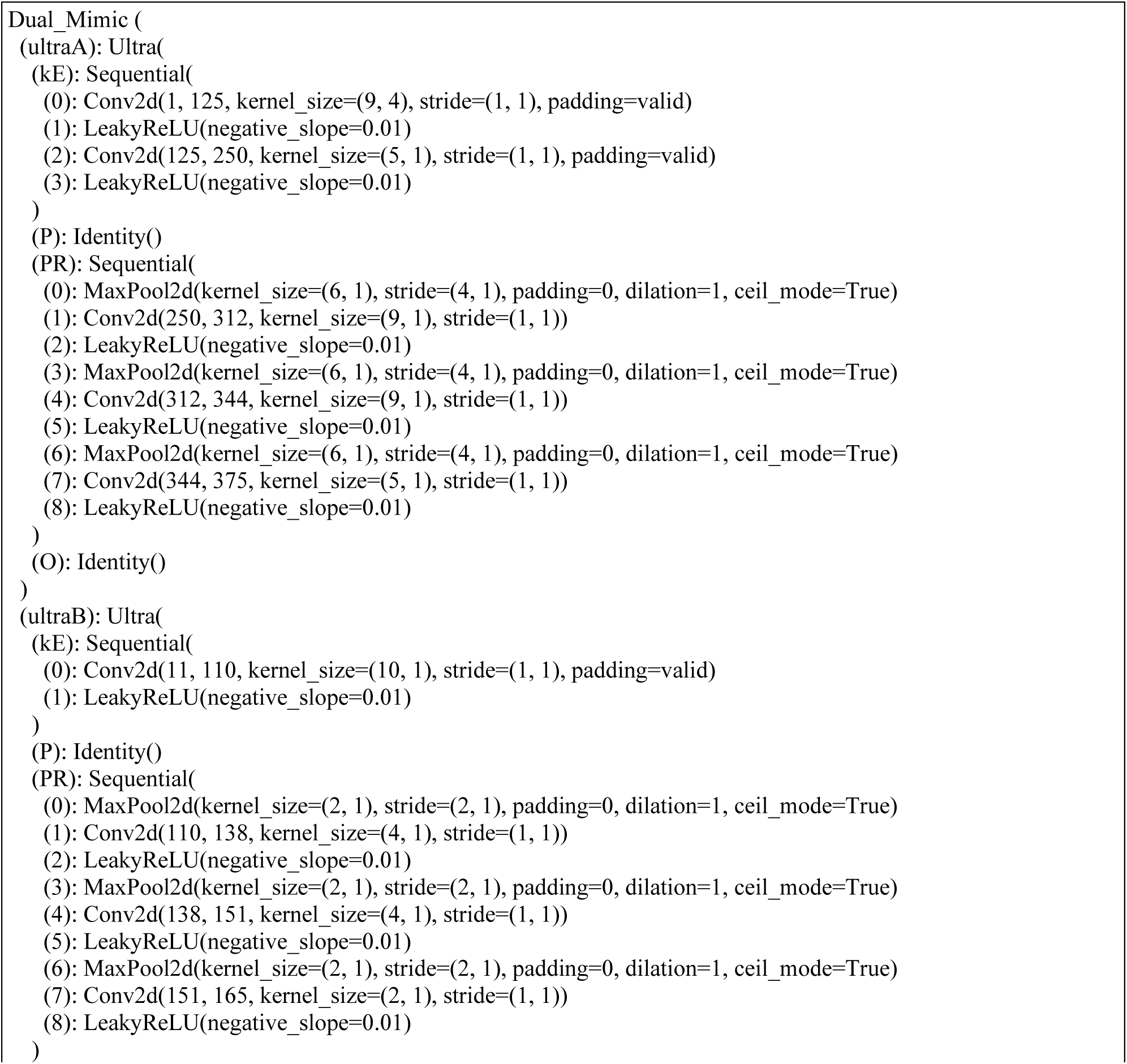
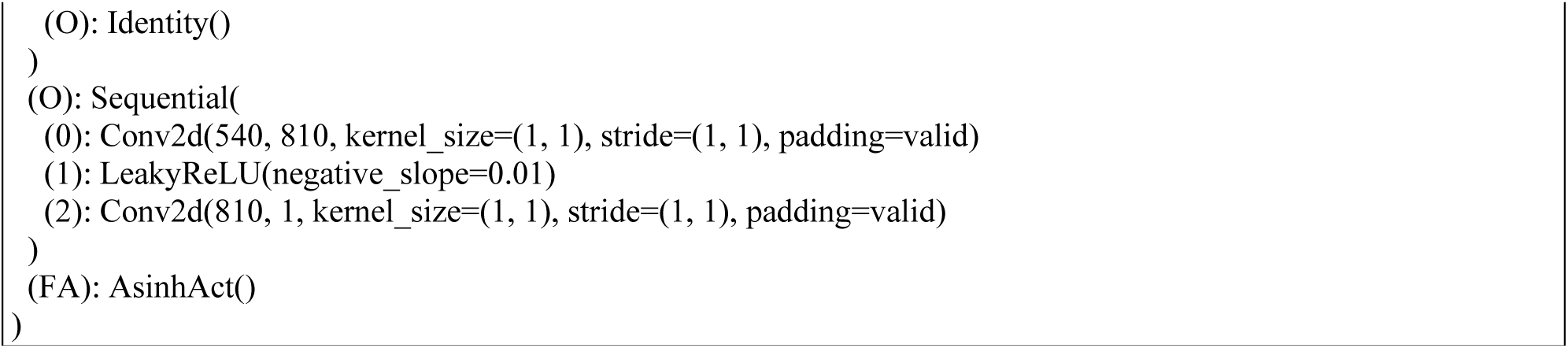
Optimized Multi-Mimic architecture for modelling CTCF-DNA interactions.

**Sup. Text. 5.**
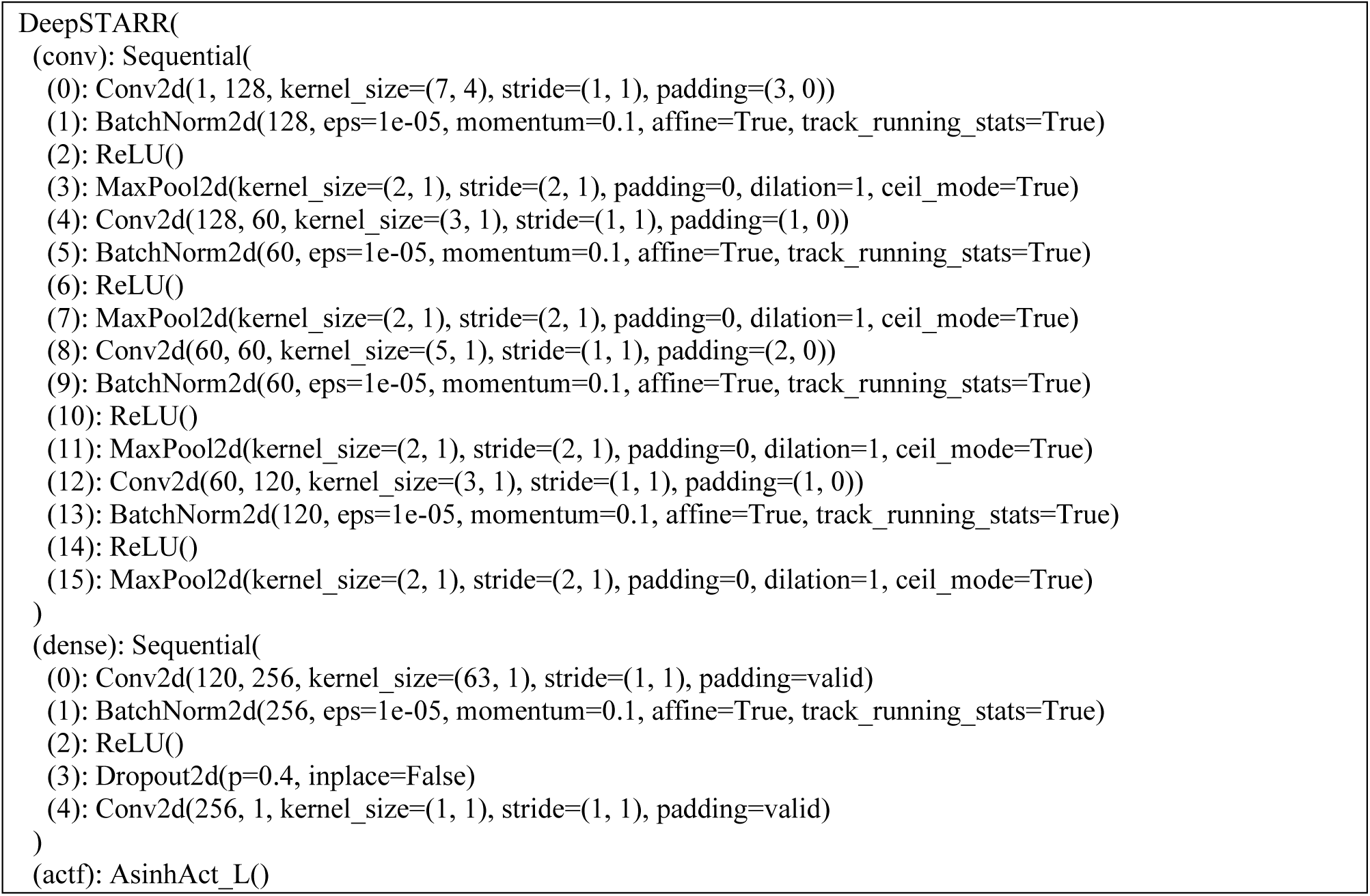
Optimized DeepSTARR architecture for modelling enhancer activity.

**Sup. Text. 6.**
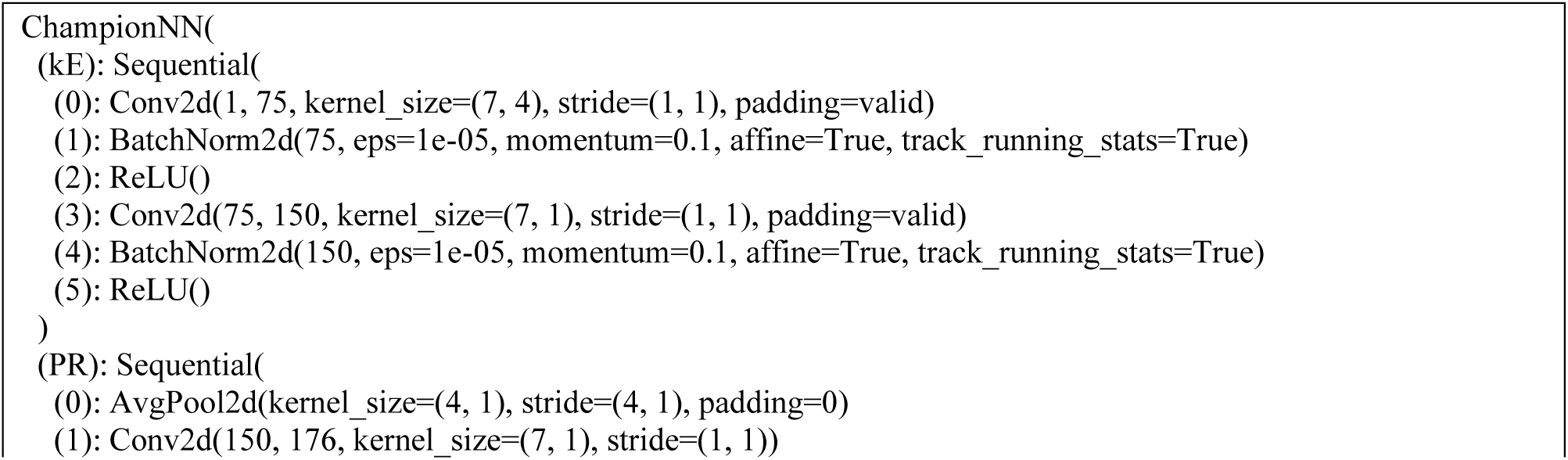
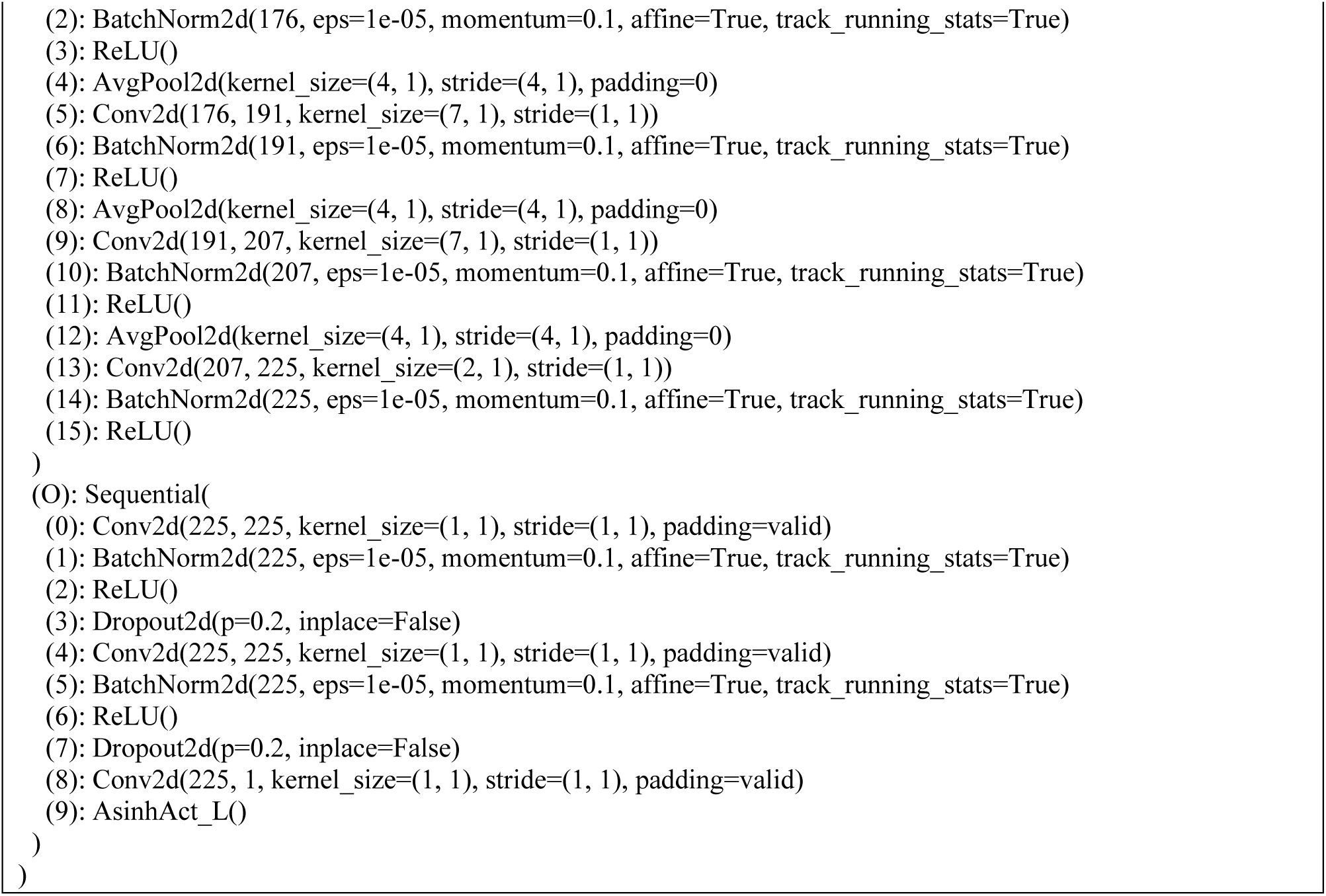
Optimized Mimic architecture for modelling enhancer activity.

## Notes

### Competing Interest Statement

The authors have declared no competing interest.

## References

Abdulnabi, H., & Westwood, J. T. (2026). Binning-based Methods for Model Fitting and Data Splitting Improve Machine Learning Imbalanced Regression. BioRkiv. 10.1101/2025.06.26.661884

Arnold, C. D., Gerlach, D., Stelzer, C., Boryń, Ł. M., Rath, M., & Stark, A. (2013). Genome-wide quantitative enhancer activity maps identified by STARR-seq. *Science (New York*, N.Y.), 339(6123), 1074–1077. 10.1126/science.1232542

Avsec, Ž., Agarwal, V., Visentin, D., Ledsam, J. R., Grabska-Barwinska, A., Taylor, K. R., Assael, Y., Jumper, J., Kohli, P., & Kelley, D. R. (2021). Effective gene expression prediction from sequence by integrating long-range interactions. Nature Methods, 18(10), 1196–1203. 10.1038/s41592-021-01252-x

Avsec, Ž., Weilert, M., Shrikumar, A., Krueger, S., Alexandari, A., Dalal, K., Fropf, R., McAnany, C., Gagneur, J., Kundaje, A., & Zeitlinger, J. (2021). Base-resolution models of transcription-factor binding reveal soft motif syntax. Nature Genetics, 53(3), 354–366. 10.1038/s41588-021-00782-6

Balcı, A. T., Ebeid, M. M., Benos, P. V, Kostka, D., & Chikina, M. (2023). An intrinsically interpretable neural network architecture for sequence-to-function learning. Bioinformatics (Oxford, England), *39*(39 Suppl 1), i413–i422. 10.1093/bioinformatics/btad271

Bartlett, M. S. (1947). The Use of Transformations. Biometrics, 3(1), 39. 10.2307/3001536

Bulyk, M. L., Johnson, P. L. F., & Church, G. M. (2002). Nucleotides of transcription factor binding sites exert interdependent effects on the binding affinities of transcription factors. Nucleic Acids Research, 30(5), 1255–1261. 10.1093/nar/30.5.1255

Chandrashekar, P. B., Chen, H., Lee, M., Ahmadinejad, N., & Liu, L. (2024). DeepCORE: An interpretable multi-view deep neural network model to detect co-operative regulatory elements. Computational and Structural Biotechnology Journal, 23, 679–687. 10.1016/j.csbj.2023.12.044

de Almeida, B. P., Reiter, F., Pagani, M., & Stark, A. (2022). DeepSTARR predicts enhancer activity from DNA sequence and enables the de novo design of synthetic enhancers. Nature Genetics, 54(5), 613–624. 10.1038/s41588-022-01048-5

ENCODE Project Consortium. (2012). An integrated encyclopedia of DNA elements in the human genome. Nature, 489(7414), 57–74. 10.1038/nature11247

Guertin, M. J., & Lis, J. T. (2010). Chromatin landscape dictates HSF binding to target DNA elements. PLoS Genetics, 6(9), e1001114. 10.1371/journal.pgen.1001114

Kelley, D. R. (2020). Cross-species regulatory sequence activity prediction. PLoS Computational Biology, 16(7), e1008050. 10.1371/journal.pcbi.1008050

Kelley, D. R., Reshef, Y. A., Bileschi, M., Belanger, D., McLean, C. Y., & Snoek, J. (2018). Sequential regulatory activity prediction across chromosomes with convolutional neural networks. Genome Research, 28(5), 739–750. 10.1101/gr.227819.117

Kelley, D. R., Snoek, J., & Rinn, J. L. (2016). Basset: learning the regulatory code of the accessible genome with deep convolutional neural networks. Genome Research, 26(7), 990–999. 10.1101/gr.200535.115

Kingma, D. P., & Ba, J. (2014). Adam: A Method for Stochastic Optimization.

Kulakovskiy, I. V, Vorontsov, I. E., Yevshin, I. S., Sharipov, R. N., Fedorova, A. D., Rumynskiy, E. I., Medvedeva, Y. A., Magana-Mora, A., Bajic, V. B., Papatsenko, D. A., Kolpakov, F. A., & Makeev, V. J. (2018). HOCOMOCO: towards a complete collection of transcription factor binding models for human and mouse via large-scale ChIP-Seq analysis. Nucleic Acids Research, *46*(D1), D252–D259. 10.1093/nar/gkx1106

Li, J., Pu, Y., Tang, J., Zou, Q., & Guo, F. (2021). DeepATT: a hybrid category attention neural network for identifying functional effects of DNA sequences. Briefings in Bioinformatics, 22(3). 10.1093/bib/bbaa159

Liu, B., Zhang, W., Zeng, X., Loza, M., Park, S.-J., & Nakai, K. (2024). TF-EPI: an interpretable enhancer-promoter interaction detection method based on Transformer. Frontiers in Genetics, 15. 10.3389/fgene.2024.1444459

Love, M. I., Huber, W., & Anders, S. (2014). Moderated estimation of fold change and dispersion for RNA-seq data with DESeq2. Genome Biology, 15(12), 550. 10.1186/s13059-014-0550-8

Maslova, A., Ramirez, R. N., Ma, K., Schmutz, H., Wang, C., Fox, C., Ng, B., Benoist, C., Mostafavi, S., & Immunological Genome Project. (2020). Deep learning of immune cell differentiation. Proceedings of the National Academy of Sciences of the United States of America, 117(41), 25655–25666. 10.1073/pnas.2011795117

McDonald, J. B., & Xu, Y. J. (1995). A generalization of the beta distribution with applications. Journal of Econometrics, 66(1–2), 133–152. 10.1016/0304-4076(94)01612-4

Min, X., Zeng, W., Chen, N., Chen, T., & Jiang, R. (2017). Chromatin accessibility prediction via convolutional long short-term memory networks with *k* -mer embedding. Bioinformatics, 33(14), i92–i101. 10.1093/bioinformatics/btx234

Phillips, J. E., & Corces, V. G. (2009). CTCF: Master Weaver of the Genome. Cell, 137(7), 1194–1211. 10.1016/j.cell.2009.06.001

Quang, D., & Xie, X. (2016). DanQ: a hybrid convolutional and recurrent deep neural network for quantifying the function of DNA sequences. Nucleic Acids Research, 44(11), e107. 10.1093/nar/gkw226

Satopaa, V., Albrecht, J., Irwin, D., & Raghavan, B. (2011). Finding a" kneedle" in a haystack: Detecting knee points in system behavior. 31st International Conference on Distributed Computing Systems Workshops, 166–171.

Singh, R., Lanchantin, J., Robins, G., & Qi, Y. (2016). DeepChrome: deep-learning for predicting gene expression from histone modifications. Bioinformatics, 32(17), i639–i648. 10.1093/bioinformatics/btw427

Singh, R., Lanchantin, J., Sekhon, A., & Qi, Y. (2017). Attend and Predict: Understanding Gene Regulation by Selective Attention on Chromatin. Advances in Neural Information Processing Systems, 30, 6785–6795.

Slattery, M., Zhou, T., Yang, L., Dantas Machado, A. C., Gordân, R., & Rohs, R. (2014). Absence of a simple code: how transcription factors read the genome. Trends in Biochemical Sciences, 39(9), 381–399. 10.1016/j.tibs.2014.07.002

Srivastava, D., Aydin, B., Mazzoni, E. O., & Mahony, S. (2021). An interpretable bimodal neural network characterizes the sequence and preexisting chromatin predictors of induced transcription factor binding. Genome Biology, 22(1), 20. 10.1186/s13059-020-02218-6

Vaswani, A., Shazeer, N., Parmar, N., Uszkoreit, J., Jones, L., Gomez, A. N., Kaiser, L., & Polosukhin, I. (2023). Attention Is All You Need.

Verheul, T. C. J., van Hijfte, L., Perenthaler, E., & Barakat, T. S. (2020). The Why of YY1: Mechanisms of Transcriptional Regulation by Yin Yang 1. Frontiers in Cell and Developmental Biology, 8, 592164. 10.3389/fcell.2020.592164

Wang, J., Zhuang, J., Iyer, S., Lin, X., Whitfield, T. W., Greven, M. C., Pierce, B. G., Dong, X., Kundaje, A., Cheng, Y., Rando, O. J., Birney, E., Myers, R. M., Noble, W. S., Snyder, M., & Weng, Z. (2012). Sequence features and chromatin structure around the genomic regions bound by 119 human transcription factors. Genome Research, 22(9), 1798–1812. 10.1101/gr.139105.112

Yang, B., Liu, F., Ren, C., Ouyang, Z., Xie, Z., Bo, X., & Shu, W. (2017). BiRen: predicting enhancers with a deep-learning-based model using the DNA sequence alone. Bioinformatics, 33(13), 1930–1936. 10.1093/bioinformatics/btx105

Zhou, J., & Troyanskaya, O. G. (2015). Predicting effects of noncoding variants with deep learning-based sequence model. Nature Methods, 12(10), 931–934. 10.1038/nmeth.3547

